# Feedback control of recurrent circuits imposes dynamical constraints on learning

**DOI:** 10.1101/2024.05.24.595772

**Authors:** Harsha Gurnani, Weixuan Liu, Bingni W. Brunton

## Abstract

Neural activity has been observed to lie on low-dimensional manifolds, constraining what new behaviors can be easily learned. We propose that beyond this geometric constraint, learning on fast timescales is limited by how neural activity can flow over time within these manifolds—i.e., by the system’s underlying dynamics. In primary motor cortex (M1), these neural dynamics are shaped not only by internal recurrence but also by sensory feedback that can continually update cortical activity. Modeling recurrent neural networks with adaptive feedback controllers in a brain-computer interface (BCI) task, we show that feedback-driven dynamics determine not just the robustness but also the flexibility of motor output. Through a control-theoretic approach, we quantitatively link learning speed and success for individual BCI decoders to the structure of input-driven network dynamics. We show that rapid learning is fundamentally limited by the network’s controllability—the ease with which inputs can steer neural activity along desired directions. Crucially, this dynamical systems perspective explains a continuous form of experimentally-observed learning variability across decoders with similar geometry, that has not been addressed previously. We also make a testable prediction that rapid adaptation to new BCI decoders depends on upstream input plasticity, such as remapping of sensory feedback, beyond local plasticity within M1. Overall, our work identifies potential network mechanisms for fast but limited motor learning, and clarifies how constraints on learning arise from both the geometry of neural activity and its underlying dynamical structure.

## Introduction

Dynamical systems frameworks have become influential in understanding neural activity structure across diverse computations including decision making, navigation, and motor control [1–12], where structured connectivity shapes the flow of neural activity over time. A critical question remains: how does this underlying dynamical structure interact with learning-related organization of neural activity, and how does it support learning of new tasks? For example, in trained recurrent neural networks, dynamical motifs can be re-used when learning a similar task [10, 13]. While latent neural dynamics have been identified in numerous neural recordings from behaving animals, it remains unclear if these dynamics play a similar role in determining subsequent learning outcomes.

To explore these questions, we were inspired by experiments in which primates learn a brain-computer interface (BCI) task. BCIs use an experimenter-defined mapping to translate recorded neural activity into motor outcomes, typically moving effectors such as a robotic limb or a cursor. One pivotal study examined how primates trained on well-calibrated BCI decoders could learn to use new decoders within a few hundred trials [14]. Crucially, not all new decoders were equally learnable; “within-manifold” decoders—those aligned with a baseline low-dimensional neural subspace (“intrinsic manifold”)—were learned more rapidly. Moreover, neural geometry in the form of this intrinsic manifold remained stable over such short-term learning. However, there was significant variability in learning outcomes, even across “within-manifold” decoders that were equally well-aligned to the low-dimensional subspace. This variability could not be understood solely through constraints on neural *geometry*. Given the growing evidence for the role of intrinsic dynamics in patterning activity in primary motor cortex (M1) [3, 4, 15–17], we hypothesized that this underlying dynamical structure would impose even stronger constraints on not just which neural “states” are accessible (e.g. those on the intrinsic manifold) but how these states can be sequenced over time. This *dynamical* constraint on entire neural trajectories could contribute to different learning outcomes for tasks with similar geometry.

To understand these dynamical constraints, we need to identify the origin of the system’s dynamics. Many previous studies have focused on primary motor cortex (M1) as a semi-autonomous dynamical system where motor preparation sets the initial conditions, after which M1 activity evolves lawfully based on its current state. Recurrent neural network models (RNNs) trained to perform motor tasks and receiving relatively simple inputs also show activity structure similar to M1 recordings [18, 19]. These results have strengthened the notion that M1 can act as a pattern generator to produce complex spatiotemporal dynamics. Despite the success of these models, they overlook observations that motor cortical activity is altered by sensory feedback, and is dependent on input from other regions such as motor thalamus [20–28]. These models also fail to recognize aspects of motor control theoretical frameworks, which propose that motor commands are continually optimized via feedback laws during movement execution [29–31]. While these control-theoretical frameworks have a rich history of interpreting and predicting behavioral dynamics, their application to understanding neural data has been more limited. Bridging this gap requires new network models that incorporate feedback [20, 32–34] and account for how neuronal interactions shape M1 activity structure for feedback control [35].

Re-conceptualizing motor cortical dynamics as being modulated by time-varying feed-back allows us to differentiate between potential mechanisms for behavioral flexibility, and the resultant constraints on learning. Animals and humans show a remarkable ability to adapt to new sensorimotor mappings on fast timescales [36–41]. If such learning causes changes to recurrent connections in motor cortex, it risks overwriting useful knowledge within an expressive pattern generator. Alternatively, learning to map sensory information into new inputs, potentially as a new context-specific map, could alter the effective behavior of the system while preserving learned structure in motor cortex. Evidence from other motor adaptation studies suggests changes upstream of M1 [42–46], such as changes in preparatory activity [47, 48] and dependence on sensory processing [41, 42, 49, 50]. Sensory feedback is also critical for successful BCI learning [51–56]. This motivated us to explore the interaction between input plasticity and preserved recurrent connectivity in generating adapted trajectories during fast BCI learning. While recent modeling studies have demonstrated similar principles for feedforward inputs that are fixed over time—contextual inputs can flexibly combine existing dynamical motifs for fast adaptation in cognitive tasks [10, 13] and generate new motor sequences [57]—, this has not yet been investigated for sensorimotor adaptation, particularly in the context of feedback control.

Motivated by the impact of feedback on motor cortical activity and the potential of inputs to reshape effective dynamics, we built feedback-driven models of recurrent circuits performing simple BCI-like tasks to understand the class of new motor computations enabled by input plasticity as well as factors that determine the success and speed of such adaptation. By modeling BCI learning phenomena [14], we show that short-term adaptation to new BCI decoders is likely facilitated by plasticity of inputs from upstream controllers, such as a remapping of sensory feedback, rather than plasticity of recurrent connections within M1. While input plasticity preserves the statistical structure of neural activity over learning, consistent with experimental observations, local recurrent plasticity not only changes intrinsic dynamics but reorganizes the covariance structure of neural activity. Crucially, we showed that beyond geometrical properties such as the alignment of BCI decoders with the intrinsic manifold, pre-existing structured dynamics affect the speed of adaptation to different decoders. We provide a quantitative account for the variability of learning outcomes across within-manifold decoders, an explanation that has been missing from related computational studies [58–62]. Rather than treating experimentally-observed variability as noise, we propose that this variability stems, at least partly, from dynamical constraints in addition to constraints on neural geometry. Lastly, by varying controller architectures, we show a dissociation between feedback controllability—a structural property of the network—and activity covariance structure during a baseline task, suggesting that low controllability of BCI decoders rather than low variance per se limits successful adaptation. We further propose the existence of control bottlenecks as an additional constraint on learning suggesting that upstream inputs to M1 during BCI use are low-dimensional. Altogether, our work identifies potential network mechanisms for short-term motor adaptation (change in control inputs) and provides a conceptual understanding (controllability and dynamical constraints) for how input plasticity shapes learning outcomes, with broad implications for studying the role of ubiquitous feedback loops—sensory and internal—in flexibly modulating diverse cortical dynamics.

## Results

### Structured dynamics in feedback-driven networks

We developed a simple computational model to study modulation of motor cortical dynamics by feedback during a BCI cursor manipulation task. We trained rate-based recurrent neural networks (RNNs) on a 2D center-out reaching task, similar to a common BCI paradigm [14] (Figure 1a). The network was cued on each trial to move the cursor to one of 8 targets. A set of readout weights (“decoder”) specified the mapping from the RNN state to the cursor’s velocity. Unlike previous computational studies that used position decoders [58–62], here we used a velocity decoder that is similar to experimental studies [14, 53, 54, 63–65], and which depends directly on the dynamical structure because cursor position is given by integration along the entire neural trajectory and not the current (final) neural state. The network received trial-specific inputs, namely the Target-cue that specified the 2D position of the trial target, and a binary Go-cue. We refer to these as “feedforward inputs”, and they were fixed within the relevant trial epochs. Another set of inputs were dependent on the position of the cursor (sensory feedback), and in some cases, network state itself; we refer to these time-varying inputs as “feedback inputs”. The mapping between sensory state and feedback inputs could be pre-specified as the 2D position error, or learned via another neural network *F* (“feedback controller”). This overall architecture with a velocity decoder thus required networks to perform a *control* task, unlike a fixed pattern generation or an input association task, and the inclusion of feedback was essential for the network to adjust its output in response to perturbations.

**Figure 1.**
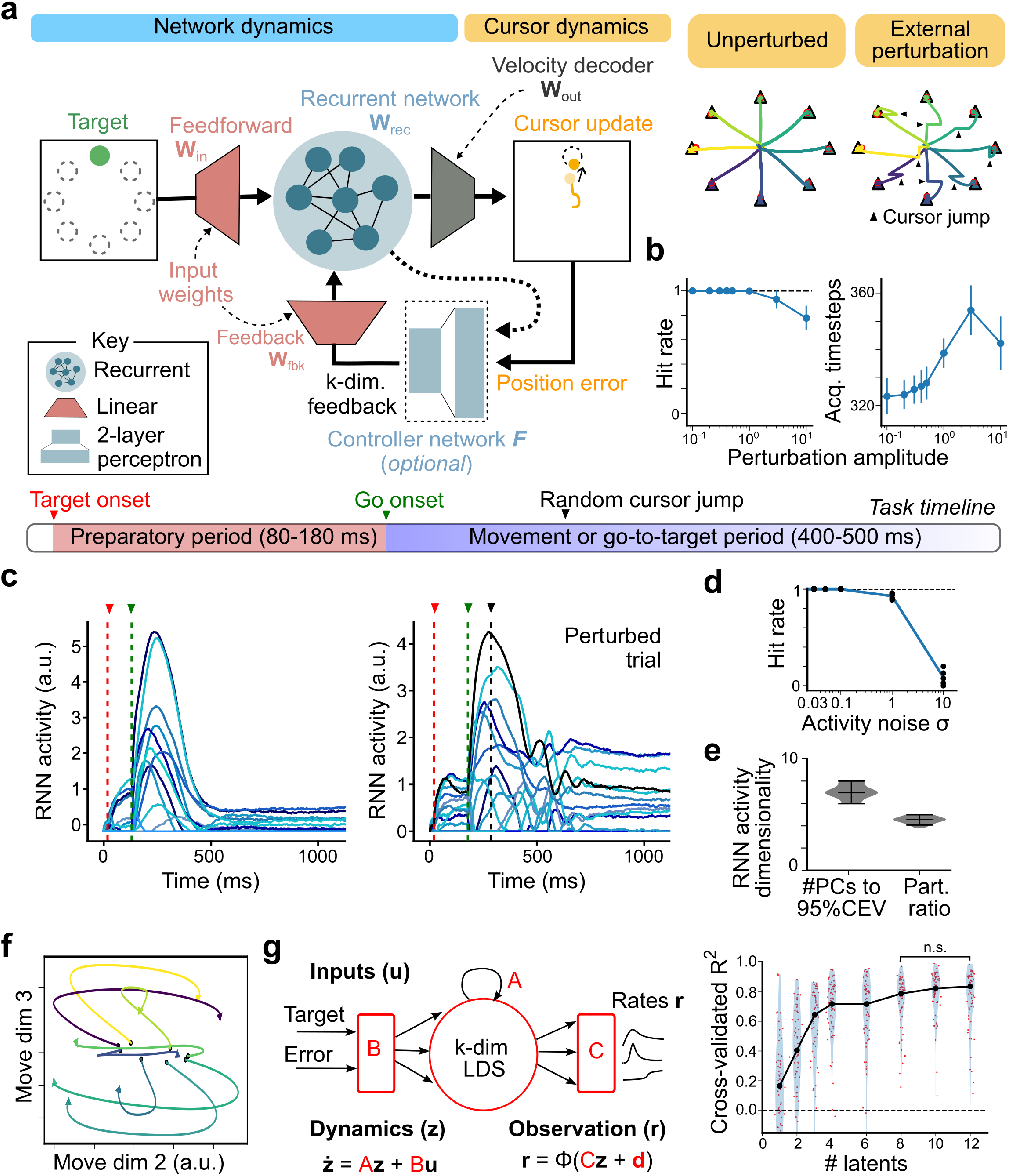
Low-dimensional dynamics in networks trained with feedback. **(a)** Network schema and behavior. (Left) Model architecture of recurrent neural network (RNN) controlling a cursor and receiving sensory feedback. The network activity **r** depends on recurrent interactions as well as two sets of inputs: feedforward and feedback. A linear readout of the network activity controls the velocity of an external cursor that is to be moved to a target location. (Right) Example cursor trajectories showing reaches to each of 8 radial targets without an external perturbation (“unperturbed”) or with an external cursor jump perturbation (example perturbations indicated by black triangles). Reaches to different targets are shown in different colors. Target locations are shown as triangles, endpoints of example reach trajectories are indicated by red circles. **(b)** Task performance in the presence of external perturbations (cursor jump), as a function of perturbation amplitude; hit rate (left) and mean acquisition time (right). **(c)** Activity of 13 example RNN units in task-trained networks; (left) an unperturbed trial with stimulus and Go-cue onset times indicated by dashed lines, and (right) a trial with cursor perturbation with the perturbation time indicated by dashed line. The two examples are reaches to different targets, with small additive activity noise (*σ* = 0.03). **(d)** Average hit rate for different levels of activity noise (n=5 networks). **(e)** Dimensionality of RNN activity measured as number of PCs to get 95% CEV and as participation ratio using the eigenvalues of the covariance matrix. Horizontal lines indicate the 5th, 50th and 95th percentile values. **(f)** Movement-related activity projected on the movement period PCs (right), colored by trial target. **(g)** A modified latent linear dynamical system was fit to RNN firing rates as observations. A low-dimensional latent dynamical system was modeled with linear dynamics *A*, which was fit along with input weights *B*, observation weights *C*, and bias terms **d**.(Right) Cross-validated *R*^2^ for models with different number of latents (k=1 to 12).

For initial training, a random decoder *W*_*out*_ was fixed while both recurrent weights *W*_*rec*_ and input weights *W*_*in*_, *W*_*fbk*_ (or the feedback controller *F*) were trained via gradient descent. To encourage networks to use feedback inputs for corrective movements, we trained them in the presence of small unintended cursor movements (see Methods). After training, we tested the networks’ robustness to larger, sudden shifts in the cursor position (Figure 1a-c). Trained networks were able to correct for these perturbations and reach the target on all trials. Secondly, while we did not add activity noise during training, we observed perfect task performance even when tested with small additive noise to the RNN activity (Figure 1d). This suggests that the trained networks were robust against both internal and external perturbations, including those not observed during training.

For low levels of noise, RNN activity was low-dimensional, with around 7-8 principal components (PCs) sufficient to capture most of the task-related variance (Figure 1e); this low-dimensional subspace corresponding to the top PCs will henceforth be referred to as the intrinsic manifold, as in [14]. RNN activity evolved in highly structured ways within this subspace; for example, we observed rotational dynamics during the movement period (Figure 1f), reminiscent of neural trajectories in primary motor cortex during reaching tasks [3]. By fitting an input-driven latent dynamical system (see Methods), we observed that low-dimensional latent dynamics reliably explained the activity structure during the task in both RNNs (Figure 1g) and BCI datasets (Figure S2). Moreover, we observed that slow recurrent processing was needed to transform error feedback into a time-varying velocity output that corrects errors over time (Figure S1, Supplementary Note), confirming that networks were not in an entirely input-dominated regime but relied on structured recurrent dynamics. These observations, along with other properties of these networks (Figure S2, S3), are qualitatively consistent with experimentally observed properties of motor cortical dynamics, enabling us to use these networks as models to identify constraints on learning in the following sections.

### Variable adaptation to different decoder perturbations

To study constraints on learning in feedback-driven networks, we adapted the paradigm of BCI decoder perturbations, as designed in [14]. After pre-training networks on the 2D BCI task with an original decoder *W*_*out*_, a new decoder *W*_*pert*_ was chosen from one of two classes of decoder perturbations – either aligned with the intrinsic manifold (within-manifold perturbation; WMP) or misaligned with it (outside-manifold perturbation; OMP) (Figure 2a; see Methods). Similar to the experiments, we ensured that selected decoders had sufficient open-loop velocities during the baseline period, as well as sufficient closed-loop velocities, so that the network produced enough variance along the new decoder to visibly move the cursor (albeit not necessarily towards the target). For experimentalists, this was needed to keep the animals motivated to perform the task, but this critical step also ensures that the behavioral impact of perturbations is similar across the two classes of decoders. Switching to these perturbed BCI velocity decoders, both WMPs and OMPs, led to a decrease in task performance as quantified by the hit rate (i.e. fraction of trials where the cursor came within 10% of the corresponding target, as defined in the original experiments). By contrast, switching to an ‘intuitive’ decoder *W*_*int*_ (see Methods), which was aligned with the intrinsic manifold and used the observed correlations between neural activity and cursor velocity, did not affect task performance.

**Figure 2.**
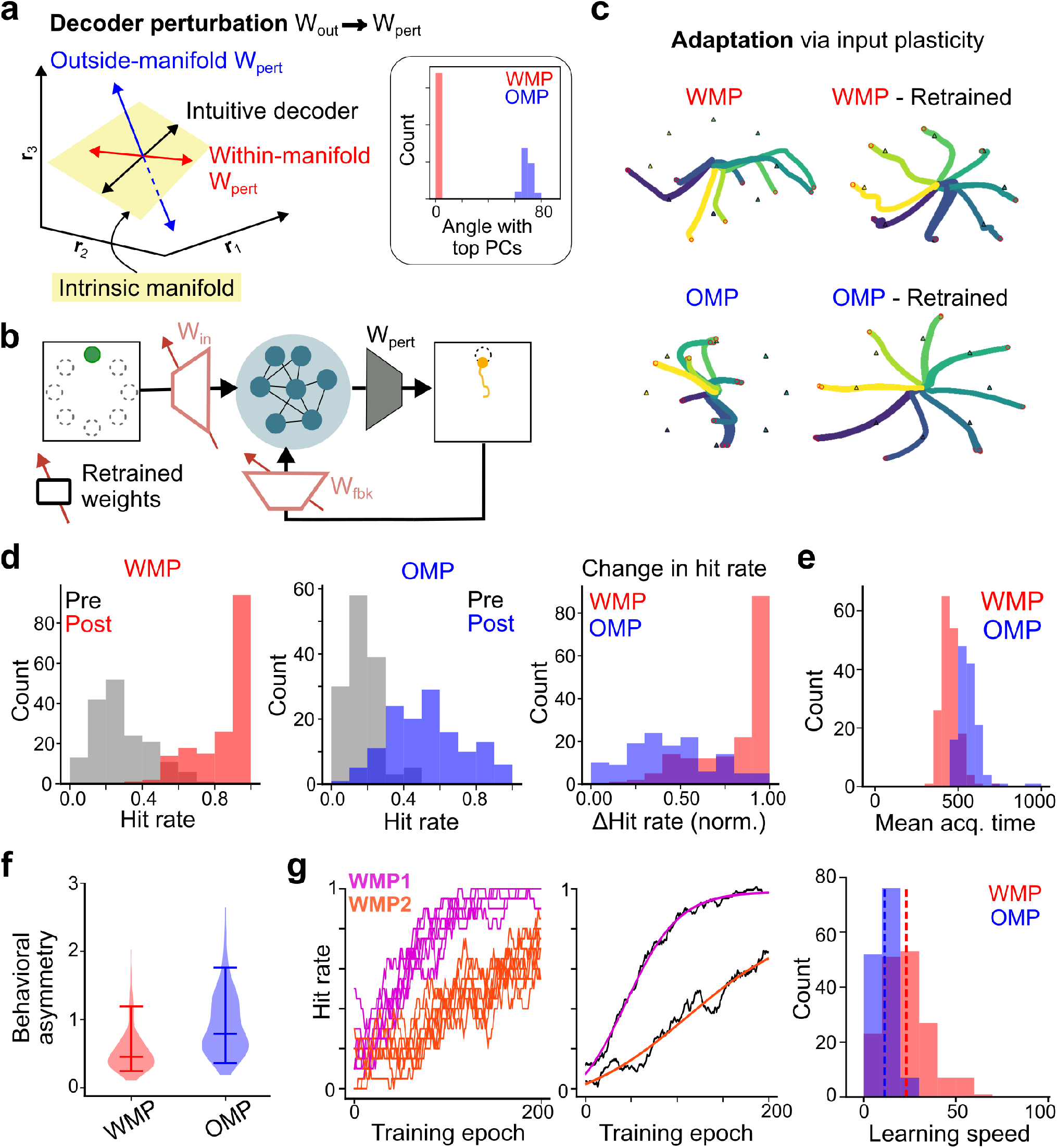
Input plasticity leads to differential learning outcomes during adaptation to decoder perturbations. **(a)** New decoders *W*_*pert*_ could be either within-manifold perturbations (WMPs, red) or outside-manifold perturbations (OMPs, blue). (Right) Distribution of angle between top PCs (intrinsic manifold) and WMPs/OMPs across different perturbations. **(b)** Model architecture with weights being retrained during adaptation to new decoders indicated by red arrow. **(c)** Example cursor trajectories after decoder perturbations (top) and after successful retraining via changes to input weights (bottom) for an example WMP and OMP. **(d)** Histogram of performance (hit rate) after decoder perturbation (grey; Pre-retraining) and after retraining for 200 trials (red/blue) for n=170 WMPs (left) and n=135 OMPs (middle). (Right) Histogram of normalized change in performance (hit rate) after 200 trials. **(e)** Distribution of mean target acquisition time for different OMPs and WMPs after retraining. **(f)** Behavioral progress asymmetry (across targets) after retraining. **(g)** (Left) Hit rate over training for 2 example WMPs (in pink and cyan). Each line is a different training iteration, with hit rate quantified as fraction of successful reaches in a 20-trial period. (Right) Average training curves for the two WMPs (black) are overlaid with a logistic fit, which is used to estimate learning speed. (Right) Distribution of learning speeds (left) across WMPs and OMPs. Dashed lines indicate median of the respective distributions.

Next, we compared how input plasticity enabled networks to adapt to new within-or outside-manifold decoders. Specifically, we retrained both feedforward and feedback input weights (see Methods) to allow the network to recover task performance, while the recurrent weights were kept fixed (Figure 2b,c). We will turn to the case of recurrent plasticity later in Figure 7. While in theory, input trajectories themselves could be directly optimized, we chose to modify the input weights to avoid such a high-dimensional optimization problem without an explicit dynamical model for the inputs. In later experiments (Figure 8), we allowed for a more expressive modification of inputs via adaptive feedback controller networks. The experimental study found that animals were able to adapt to within-manifold decoders more easily than outside-manifold decoders [14]. In our networks, we similarly noted that recovery of task performance varied across different perturbations. There were both within-class (i.e. within WMPs and within OMPs) and across-class (i.e. WMP vs OMP) differences for improvements in task performance, but with greater learning in WMPs on average (Figure 2d; median normalized improvement in hit rate: WMP=0.9, OMP=0.4). We also observed that average acquisition times (i.e. time to reach the target on each trial) were higher for OMPs (Figure 2e; target acquisition time, median [90% CI]: WMP=441 steps [369,526], OMP=554 steps [485,666]). While within-manifold decoders showed greater recovery with learning, task performance was not similar for all targets. We quantified behavioral asymmetry as the variation in average speeds (progress) towards different targets for the same decoder, and observed that this asymmetry was significantly higher than zero (Figure 2f), as was recently described in experimental data as well [62]. Overall, these behavioral differences for learning of new decoders are consistent with experimentally observed variability in learning outcomes in primate BCI experiments [14].

While all perturbations were retrained for a fixed number of iterations, we observed that learning curves (hit rate across training epochs) were highly consistent for individual perturbations across different training runs (different sequence of stimuli), but that they were different across perturbations, even when both decoders were “within-manifold” (Figure 2g). On average, WMPs had higher learning speeds than OMPs but showed substantial variability (learning speed, median [90% CI]: WMP=24 [8,50], n=170, OMP=12 [6,20], n=135, p=2*e* − 7; Mann-Whitney U test). Thus, beyond categorical differences between OMPs and WMPs, there were also differences in the amount and speed of task performance recovery across within-manifold decoders, both in our models and in experimental data (see [14], Figure 2 and Extended Data Figure 2).

### Input plasticity does not alter statistical structure of neural activity

What changes in neural activity support this behavioral learning? Previous studies have characterized that within-session learning leads to relatively little change to neural repertoires i.e. the distribution of population activity states [66]. They observed that even after successful behavioral adaptation, neural activity remained within the same intrinsic manifold although individual target-specific neural activity patterns had shifted within this manifold. Thus, for input plasticity to be a potential mechanism for experimentally-observed short-term learning, it must preserve the statistical structure of neural activity. In networks retrained with input plasticity, the covariance structure of neural activity, pooled across all targets, looked largely similar before and after adaptation to new BCI decoders (Figure 3a), for both within- and outside-manifold decoders. We confirmed that most of the neural variance was restricted to the original intrinsic manifold for both OMPs and WMPs (Figure 3b), and that with input plasticity, the change in fractional variance along the original top 8 PCs was small (ΔFractional variance, median [90%CI]: WMP=-0.03[-0.10,0.01], OMP=-0.07 [-0.13,0.06], Figure 3c). We also observed that the relative distribution of variance along the original PCs was similar before and after learning (covariance structure similarity, median [90% CI]: WMP=0.97 [0.82, 0.99], OMP=0.94 [0.80,0.98]) (Figure 3e,f). While the activity manifold remained similar, changes in activity levels could still lead to altered absolute variance along the output direction (decoder). Indeed, we observed an increase for variance along the outside-manifold decoders (Ratio of post-to-pretraining variance along decoder: WMP=0.9, OMP=2.7, p<1*e*-7, MWU) (Figure 3d). By contrast, variance along the within-manifold decoders and the resultant distributions of cursor speeds remained relatively unchanged, consistent with experimental observations [66]. Thus, input plasticity largely preserves the statistical structure of neural activity (Figure 3g) while enabling new behavioral trajectories.

**Figure 3.**
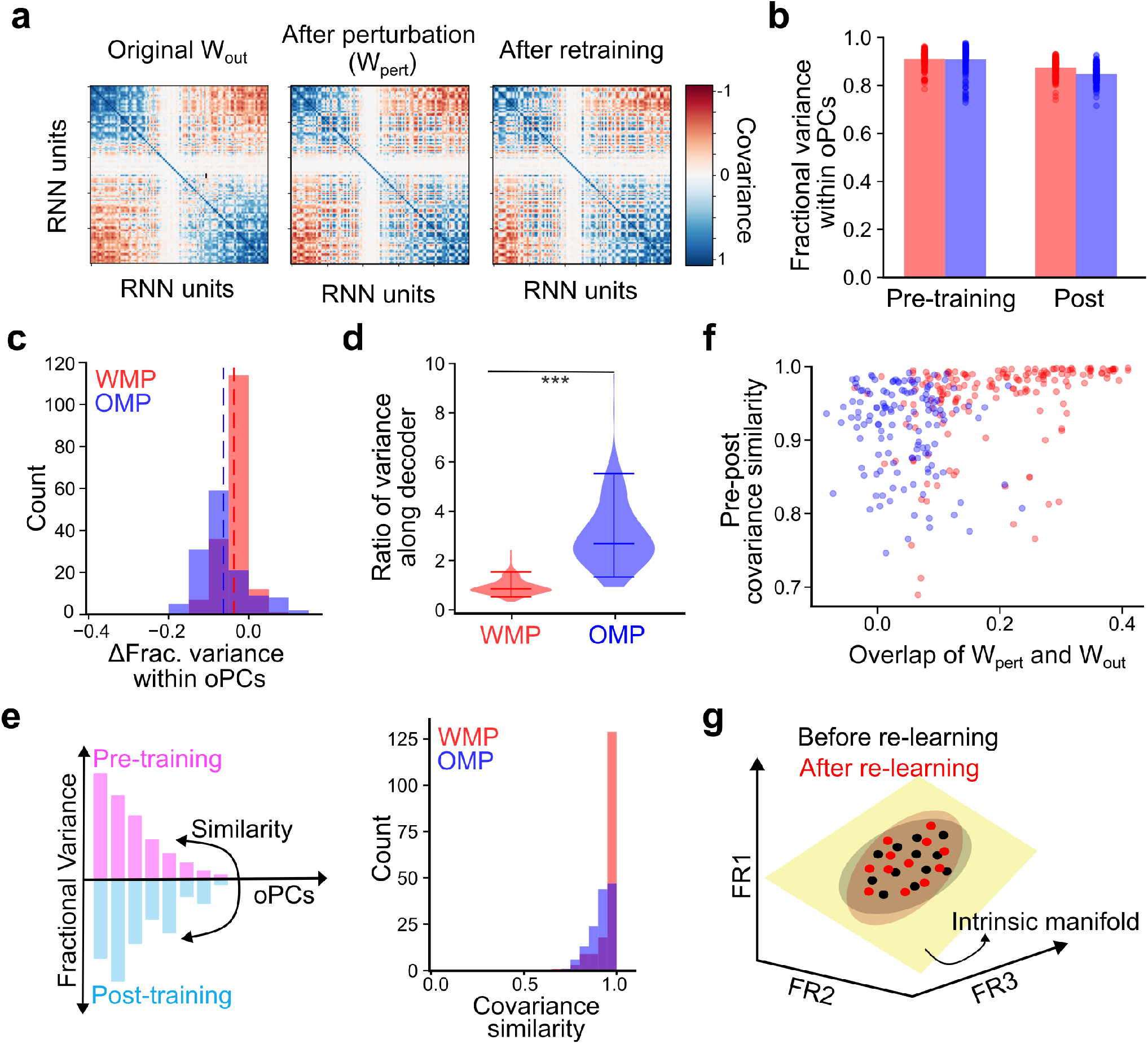
Statistical structure is largely conserved under input plasticity. **(a)** Activity covariance structure for an example network, for the baseline task (left), after introducing the new perturbed decoder *W*_*pert*_ (middle), and after retraining input weights (right). **(b)** Fractional variance within original intrinsic manifold (n=8 top PCS) for different WMPs (red, n=170) and OMPs (blue, n=135), before and after retraining. Scatter denotes individual decoder perturbations, bars indicate median across perturbations. **(c)** Distributions of change in fractional variance within original top PCs. Dashed lines indicate the medians. **(d)** Ratio of post- to pre-retraining activity variance along the perturbed decoder, shown as a distribution. Horizontal lines indicate the 5th, 50th and 95th percentile values. **(e)** Similarity of covariance structure was measured as the overlap between vectors describing the relative variance along different original PCs (oPCs). (Right) Distribution of covariance similarity for different WMPs and OMPs. **(f)** Covariance similarity versus similarity of perturbed and original decoder. **(g)** Summary: Activity remains within the intrinsic manifold after retraining, with similar covariance structure.

### Dynamical constraints on learning

While alignment with the intrinsic manifold, a *geometric* property, has been proposed as the determinant of learning differences between WMPs and OMPs [14], it is insufficient to explain all constraints on learning, in particular the variability across WMPs (all WMPs lie within the intrinsic manifold). Previous analyses did not find a systematic relationship between learning outcomes for WMPs and various geometric properties [14]. We instead propose that understanding this variability requires a focus on *dynamical* properties.

As an illustrative example, Figure 4 shows different neural trajectories that might be required to produce desired behavioral outcomes. In a network with its own internal recurrent dynamics, different trajectories can be flexibly generated using a combination of input and recurrent drive (as well as by setting appropriate initial conditions), with inputs provided by a controller network via input weights *B*1 and *B*2 (Figure 4a). *Controllability* is an important property of the network which determines the ease of moving network activity from one state in any particular direction. Moving along the recurrent flowfield or in directions coupled to large input weights requires small controller output, i.e. it has a lower control cost. Figure 4b shows three example trajectories (*T* 1,*T* 2,*T* 3) that are within the same two-dimensional stable subspace (same variance subspace) and that consist of “within-manifold” activity states. Yet, they are not equivalent. Trajectory *T* 1 is mostly aligned with the recurrent flowfield and requires little input drive, while Trajectory *T* 3 requires stronger, potentially infeasible input drive to go against the intrinsic (recurrent) flowfield. Trajectory *T* 2 also requires stronger input drive than *T* 1 as the neural state first needs to be pushed out into a less controllable direction (as determined by the weaker input weight *B*2). These *dynamical* properties allow us to begin differentiating between within-manifold *trajectories*. The last trajectory *T* 4 is partly outside the low-dimensional dynamical subspace; if the inputs and recurrent dynamics are entirely confined to the 2D subspace, then no amount of control input can produce such a trajectory. Further, if the recurrent flowfield remains fixed, then reducing the control costs (e.g., if there are upper bounds or metabolic costs on inputs) or mapping constrained external inputs via new input weights over learning requires input weights *B*1 and *B*2 to be modified to different degrees. For example, *T* 2 may be learned via a small rescaling of *B*2 whereas *T* 3 may require significant rotation and rescaling of input weights. Can we define difficult within-manifold perturbations, or difficult decoders in general, as those less aligned to controllable directions, or where the necessary dynamics to produce behaviorally-appropriate neural trajectories are significantly different from existing flowfields?

**Figure 4.**
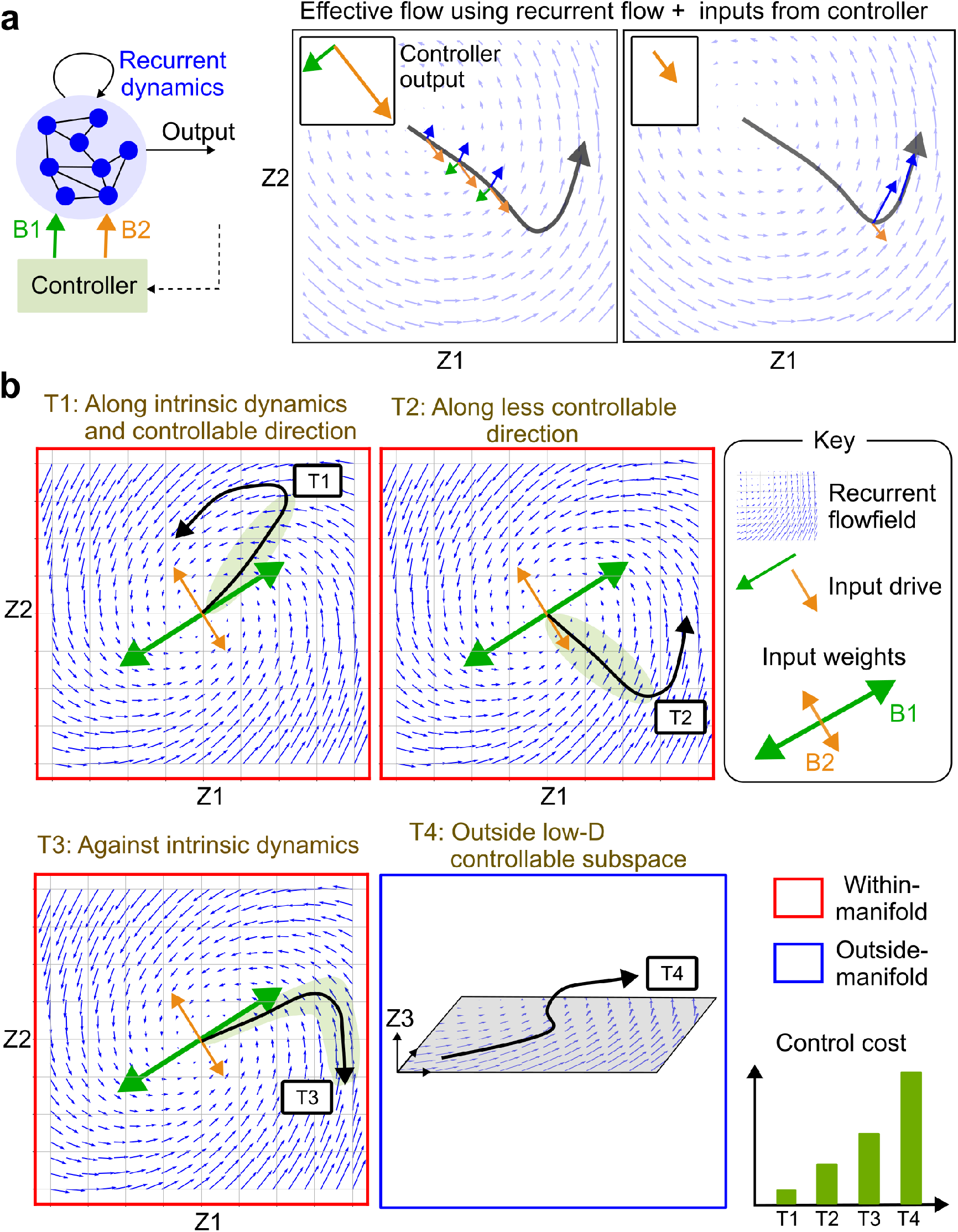
Dynamical constraints in controlled networks. **(a)** Neural activity over time is shaped by both recurrent dynamics and control inputs provided via input weights *B*1 and *B*2. (Right) An example neural trajectory with net contribution of recurrent flow (blue), and control inputs via *B*1 (green) and *B*2 (orange). The corresponding controller output, and hence the control cost, varies across parts of the trajectory as differences in controllability arise due to recurrent dynamics and unequal input weights. **(b)** Trajectories T1-T4 differ in control costs due to varying alignment with intrinsic dynamics and controllable directions. (The parts of the trajectory that require control inputs are highlighted by green ellipses.) Input weights B1 are larger than B2, creating more and less controllable directions. Trajectory T1 requires minimal input as it is aligned with highly controllable directions whereas T2 requires larger inputs to push along a less controllable direction. T3 requires strong external control inputs to overcome the recurrent flow field. T4 may be infeasible, or require even greater inputs, if it is poorly aligned with the low-dimensional dynamics/controllable subspace.

To answer this, we first quantified the local controllability spectrum for the nonlinear dynamics of our pretrained networks at multiple points within the neural state space (see Methods), and observed that it was dominated by a few highly controllable modes (Figure 5a). This means that at any given part of the neural state space, there are only a few directions along which the state can be moved easily i.e. using small inputs in addition to the recurrent drive. Next, for each new decoder *W*_*pert*_, we quantified the average overlap between the new within-manifold or outside-manifold decoder with the most controllable subspace at each state, typically a 10 dimensional subspace (Figure 5b). We observed that while controllability was lower for both perturbations compared to the intuitive decoder, OMPs had much lower overlap with the controllable subspace, and there was some variability across WMPs (Overlap of *W*_*pert*_ with feedback-controllable subspace, normalized by overlap of intuitive decoder: WMP=0.71 [0.47,0.86], n=170, OMP=0.29 [0.20,0.45], n=135).

**Figure 5.**
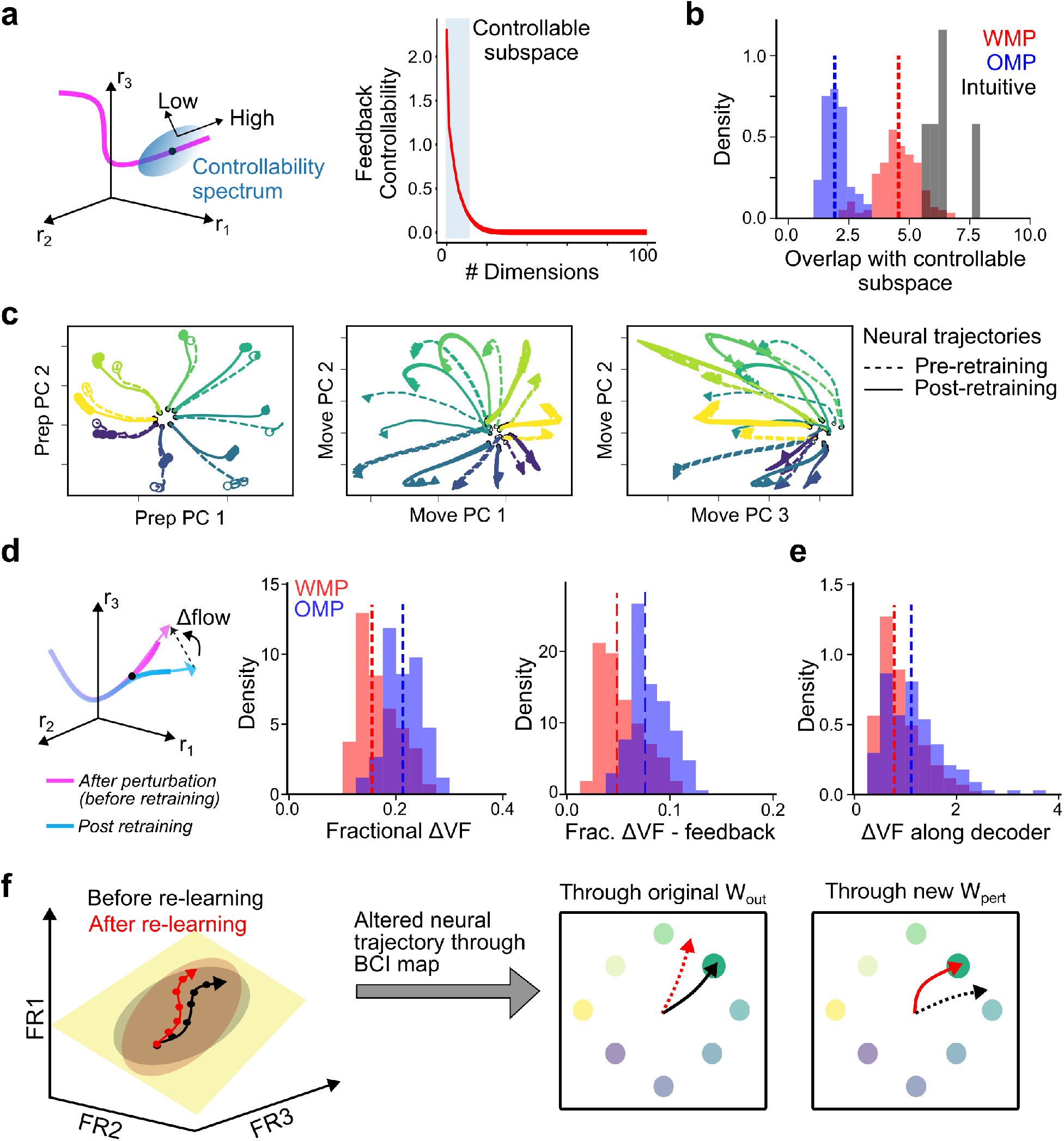
Dynamical features of adaptation to decoder perturbations. **(a)** The controllability spectrum can be calculated locally around any point in neural state space, which defines highly or poorly controllable directions. (Right) Average feedback controllability spectrum across different parts of state space and n=5 networks. The steep falloff suggests only a few directions are highly controllable locally around any point. **(b)** Overlap of the most controllable directions with the decoder (WMP or OMP *W*_*pert*_ and intuitive decoder) for n=170 WMPs, n=135 OMPs, n=5 intuitive decoders. **(c-e)** Retraining leads to a change in flow fields (or vector-field VF), supporting a small dynamical reorganization to produce new trajectories. **(c)** Target-specific neural trajectories for an example network, after introducing new decoder *W*_*pert*_ (pre-retraining; dashed), and after adapting input weights (post-retraining; solid). Trajectories are visualized in the space of top preparatory (prep) and movement (move) period principal components. **(d)** Distribution across WMPs and OMPs, of the mean change in the total flow (left) and the feedback-driven flow component (right), normalized by the total flow at each point in state space, and averaged across observed neural states during post-retraining task performance. **(e)** Normalized change in flow along the perturbed decoder direction is much higher as new behavioral trajectories are produced with successful adaptation. **(f)** Summary: Changes in effective input-driven dynamics produces new neural trajectories within the same intrinsic manifold. Altered neural trajectory, passed through the new BCI decoder, restores desired behavior.

To assess the suitability of pre-existing flowfields, we examined the temporal evolution of neural activity pre- and post-learning and directly quantified changes in the flowfield at various points in neural state space. We focused on local flowfield changes rather than fixed point structure, as the network can successfully perform the task by briefly passing through the target (without requiring the re-emergence of appropriate fixed points to stabilize the cursor at the targets). While RNN activity remained within the original manifold, the target-specific neural trajectories were altered (Figure 5c). Although both recurrent interactions and inputs shape the net flowfield (Figure 4a), here the dynamical reorganization was due to changes in the input-driven component. We separately examined both the total change as well as feedforward- and feedback-driven components of flowfield changes (Figure 5d). We observed that these changes were small (~ 10-30% of total original flow), and that the feedback-driven component of this change was different between WMPs and OMPs (Fractional ΔVF, mean±SD: WMP=0.15 ±0.04, OMP=0.21 ±0.03; Frac. ΔVF-feedback: WMP=0.05 ±0.02, OMP=0.08 ±0.02, p<1*e*-7, MWU). While these changes in flowfields are expected to be small when the recurrent component still dominates the effective flowfield, they can add up across timesteps to produce more varied trajectories. The proportional reorganization of flow along the readout was larger, for both WMPs and OMPs (Frac. ΔVF along decoder: WMP=0.90 ±0.42, OMP=1.19 ±0.66, Figure 5e), as expected from changes in cursor trajectories over learning. Similar behavioral and neural outcomes were also observed using a local learning rule (Figure S4). Thus, in addition to the misalignment with the intrinsic manifold, outside-manifold perturbations were also characterized by lower controllability and requiring greater dynamical reorganization. Further, the small changes in the statistical structure of neural activity (Figure 3) suggest that learning occurred by a re-purposing of activity patterns from the same neural manifold but using new inputs for re-associating them with new reach targets and producing them as a part of new temporal sequences (Figure 5f).

### Learning speed depends on feedback-driven dynamics

We next turn to the question of whether these dynamical properties explained variability in learning outcomes. We observed that both metrics of adaptability — learning speed and the improvement in hit rate — co-varied with changes in feedback-driven flowfields (Figure 6a,b). A linear mixed-effects model with two predictors—feedback controllability of the perturbed decoder and feedback-driven flowfield changes—could explain a large fraction of the learning variability (Figure 6c,d) (FBK model, prediction of learning speed, cross-validated *R*^2^=0.52 [0.35,0.63]; prediction of normalized ΔHit rate, *R*^2^=0.69 [0.45,0.79]). Importantly, these predictors were significant for explaining within-class variability beyond the mean differences between WMPs and OMPs captured by the group-specific intercept (Figure 6a,b, Figure S5).

**Figure 6.**
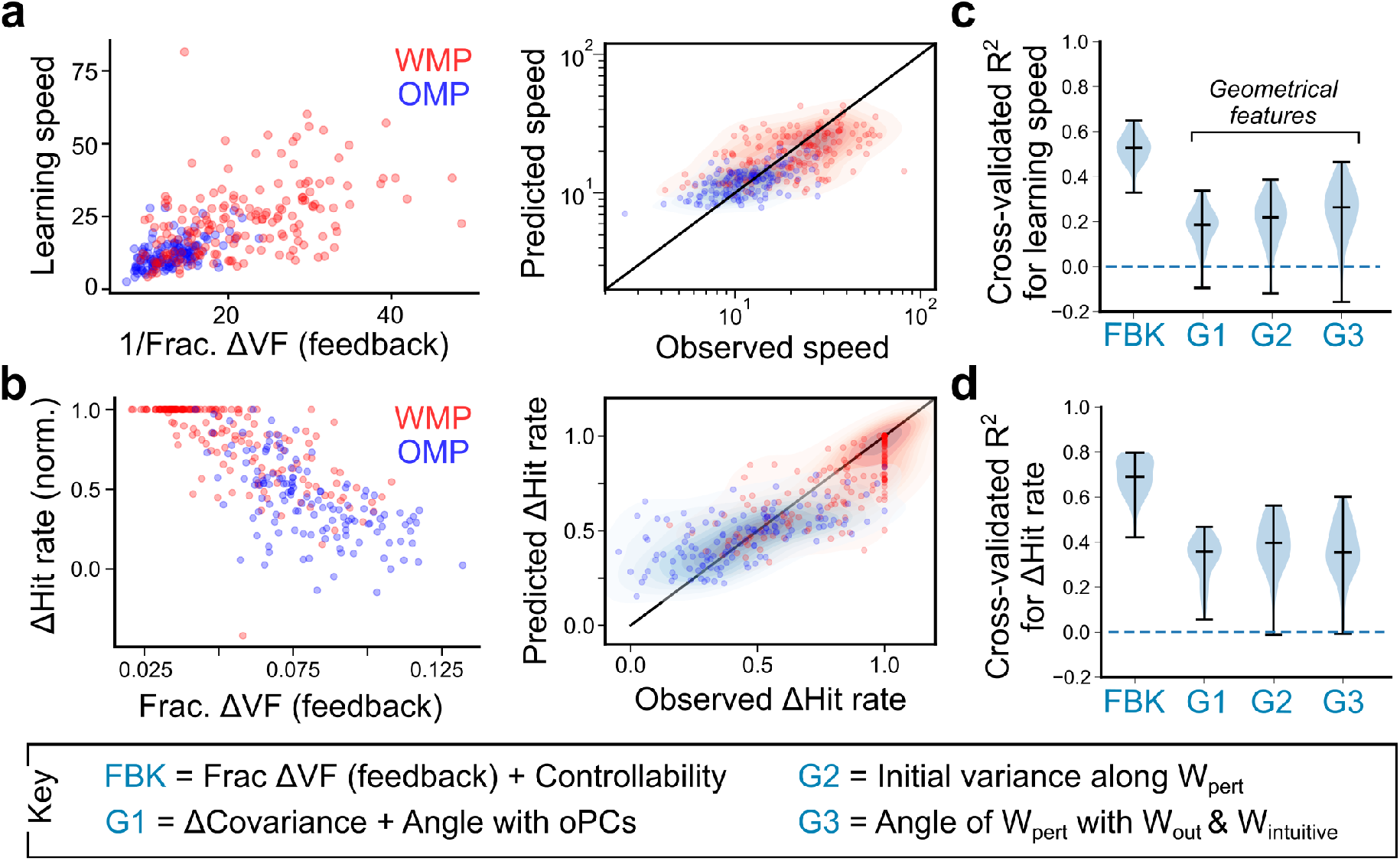
Dynamical constraints explain learning variability. **(a)** (Left) Learning speed versus change in feedback-driven flow, for WMPs (red, n=170) and OMPs (blue, n=135). (Right) Learning speeds predicted using the FBK model, against the true speeds. FBK model includes feedback controllability and changes in feedback-driven flow as predictors, as well as a group-specific (WMP/OMP) intercept. Scatter denote individual decoder perturbations, and shaded areas show different quantiles. **(b)** Similar to (a), but for normalized change in hit rate. **(c-d)** Cross-validated *R*^2^ (n=40 iterations) for 4 different models, predicting learning speeds in (c) and normalized change in hit rate in (d). Horizontal lines show extrema and median values of the distribution. All models include 2 predictors along with a group-specific intercept (fixed effect) that captures categorical differences between OMPs and WMPs. Models G1, G2 and G3 use different geometrical predictors; see Methods for details. Also see Figure S5.

This predictability was stronger for changes in feedback-driven flow as compared to feedforward-driven flow or feedforward controllability, even though learning was poorer when only feedback weights were retrained (normalized ΔHit rate with fixed *W*_*inp*_, WMP = 0.22 [-0.06, 0.66], OMP=0.16 [-0.13,0.42], n=55, p=0.09). This may be because changing the target-specific inputs (bias) is an essential and indeed larger component of the total change in effective flow, for both within- and outside-manifold perturbations. Regression models using previously characterized geometrical features such as changes in variance within the intrinsic manifold (*G*1), initial variance along the new decoder (*G*2), or angle between initial and perturbed decoders (*G*3), were all poorer at predicting learning speed (median *R*^2^=0.19, 0.22, 0.26 respectively) as well as the change in hit rate (*R*^2^=0.36, 0.41, 0.35 respectively). Importantly, these geometrical features were less significant in explaining additional variability beyond the mean differences (Figure S5), and including these predictors in the FBK model did not provide additional explanatory power. Interestingly, neural geometry or initial behavioral impairment did not explain experimental WMP variability either (Extended Data Figure 6 in [14]). Our computational modeling shows that controllability along the new decoder and changes in input-driven flow shaped learning variability across different decoder perturbations and constitute *dynamical* constraints on learning.

### Recurrent plasticity does not lead to variable learning outcomes

While we have focused on input plasticity as the mechanism underlying short-term adaptation, we also investigated the plausibility of recurrent plasticity as an alternative. Previous studies with updates of recurrent weights observed that all perturbed decoders were learnable, but differences between within-manifold (WMP) and outside-manifold perturbations (OMPs) could be attributed to (i) imperfect error feedback [58], (ii) larger synaptic changes for OMPs [59], or (iii) smaller gradients for OMPS [60, 61]. In this study, we examined the consequence of learning via recurrent plasticity using the noisy gradient estimates in trial-to-trial learning (Figure 7a, Figure S6). Similar to previous modelling work but unlike experiments [14], we observed nearly perfect recovery of task performance within the tested duration and no differences between WMPs and OMPs (Figure 7b), even for slower learning rates (Figure S7). We also observed that the trajectories were straighter after relearning, and with little behavioral asymmetry across targets for both OMPs and WMPs. Further, we observed that recurrent plasticity led to greater reorganization of neural activity structure, with reduction in fractional variance along the original PCs (Figure 7c) and realignment of activity to increase variance along the new decoder, particularly for OMPs. Moreover, the underlying dynamics were altered to a much greater extent (Figure 7d). These results suggest that neural reorganization under recurrent plasticity is inconsistent with experimental observations of relatively stable neural geometry. Recent data-driven modeling of the original experimental data also found that upstream rather than local recurrent plasticity better explains neural activity structure over short-term learning [67].

**Figure 7.**
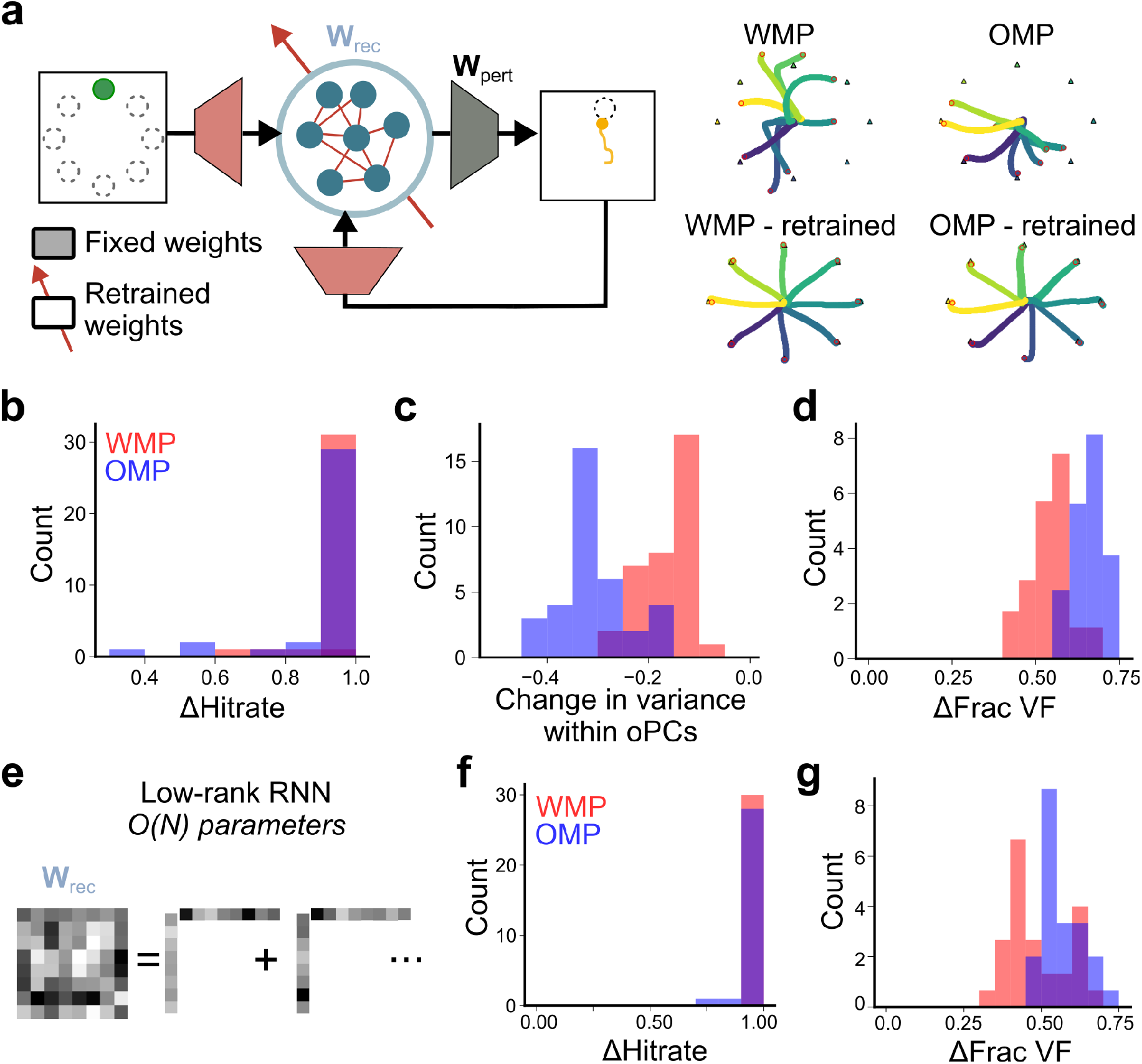
Recurrent plasticity does not lead to variability in learning outcomes. **(a-d)** Learning outcomes of recurrent plasticity in full-rank RNNs. **(a)** (Left) Schematic showing retraining of recurrent weights. (Right) Cursor trajectories after decoder perturbation (top) and retraining recurrent weights (bottom). **(b)** Distribution of normalized change in hit rate, **(c)** change in variance within original PCs and **(d)** change in recurrent flowfields, for n=35 WMPs (red) and n=35 OMPs (blue). **(e-g)** Learning outcomes of recurrent plasticity in low-rank RNNs (rank=4). **(f)** Distribution of normalized change in hit rate and **(g)** change in recurrent flowfields, for n=30 WMPs (red) and n=30 OMPs (blue). Also see Figure S7.

To assess whether better learning may be attributable to the larger number of modifiable parameters or a greater expressivity of the dynamics under recurrent plasticity, we examined adaptation to new decoders in pretrained low-rank networks [68]. By fixing recurrent weights to be constructed as a sum of rank-1 components, we ensured that the number of modifiable parameters in plastic low-rank networks was comparable to that of input plasticity (800 vs 700, Figure 7e). Despite the lower number of parameters, both OMPs and WMPs were equally learnable and accompanied by significant reorganization of the effective flowfields (Figure 7f,g). Altogether, this suggests that recurrent plasticity is unlikely to be the main driver of within-session adaptation observed in [14] as the lack of behavioral variability and the large degree of neural reorganization are both inconsistent with experimentally-observed learning [14, 66].

### Control bottlenecks shape constraints on learning

Up until now, the feedback input was a simple remapping of a two-dimensional error signal. How does the expressivity of the feedback mapping and the dimensionality of feedback inputs influence learning outcomes? To examine this, we introduced a controller network *F* that produced a *k*-dimensional output, either via feedforward processing in a 2-layer network or via recurrent processing (i.e. the controller network’s state determined the feedback). This expressivity allows for changes in areas upstream of motor cortex, such as cerebellum, parietal and premotor cortices. The *k*-dimensional control signal was provided via a set of fixed projection weights to the main recurrent network (whose activity is read out by the decoder), while the network *F* was kept plastic for adaptation (see Methods). The control signal was set to be either low-dimensional (*k* = 4, Figure 8a-c), which defined a control “bottleneck”, or it could be higher dimensional (*k* = 12, Figure 8d). We could also change the number of modifiable parameters by changing the dimensionality of the hidden layer (Figure 8b) or using a recurrent controller (Figure 8c) without changing the “bottleneck” determined by the output dimensionality of network *F*. For all controllers with the 2-layer feedforward architecture, we provided a 10-D projection of the main recurrent network’s state as input in addition to the error signal, so that a flexible, state-dependent policy could be learned. The recurrent controller instead could use its own state dependence on the error signal’s history to learn a flexible feedback policy.

**Figure 8.**
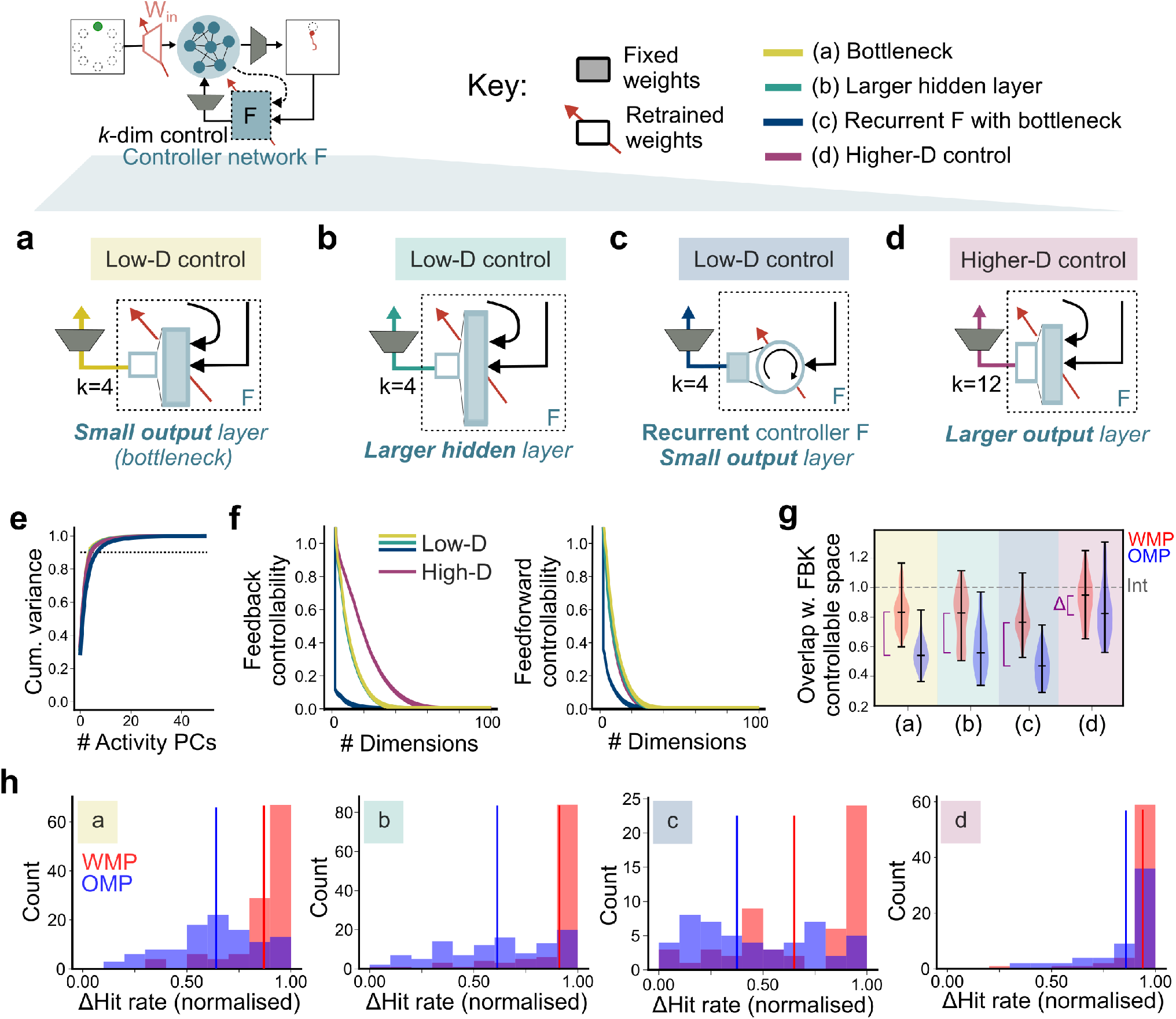
Learning outcomes depend on dimensionality of control input. **(a-d)** Networks with different controller architectures, where adaptation to decoder perturbations occurs via plasticity of input weights and controller network *F*. Controller networks in (a)-(c) have low-dimensional output, whereas controller network in (d) has a higher-dimensional output. Two-layer feedforward controllers in (a), (b) and (d) sample both the cursor position error and the RNN state, process these in a hidden layer, and have different number of output units (4 or 12); learning occurs at the weights from the hidden to the output layer of *F*. In (c), networks have a recurrent controller, with 50 hidden units and a 4-dimensional output layer, and learning occurs at the recurrent weights of *F*. For all architectures, the projection weights *W*_*fbk*_ that transform the *k*-dimensional controller output into input currents for the main RNN are fixed. **(e)** Cumulative variance for varying number of activity principal components (averaged across n=3 networks of each architecture). Color indicates different architectures from (a)-(d), as indicated in the key. **(f)** Feedback (left) and feedforward (right) controllability spectra for network architectures in (a)-(d), averaged across n=3 networks each. **(g)** Distribution of overlap of different within-manifold (WMP, red) and outside-manifold (OMP, blue) decoders with the feedback controllable subspace, normalized by the overlap of the intuitive decoder (Int). The difference between WMPs and OMPs within each type of network is highlighted as Δ. **(h)** Amount of learning for within-manifold (WMP) and outside-manifold (OMP) decoders for networks with different controller architectures, as shown in (a)-(d). For networks in (d) with higher-dimensional controller output, both WMPs and OMPs have good learning outcomes.

While these networks had similarly low-dimensional activity during the baseline task (Figure 8e), they had different feedback controllability spectra. As expected, networks with a control bottleneck had fewer highly controllable modes whereas networks with more independent input channels had an expanded controllable subspace, with a slowly decaying controllability spectrum (Figure 8f). We also observed that the difference in decoder overlap with the feedback-controllable subspace for outside-manifold versus within-manifold decoders was smallest for the architecture with higher-dimensional feedback (Figure 8g; ΔOverlap (WMP − OMP): (a) 0.29, p<e-26, (b) 0.27, p<e-5, (c) 0.29, p<e-14, (d) 0.12, p=0.004). Moreover, the overlap with the feedback controllable subspace was, on average, higher for both kinds of decoders—WMPs and OMPs—in this architecture with higher-dimensional feedback. By contrast, the geometric alignment of OMPs with the initial task-specific intrinsic manifold (high variance PCs) was similarly low across all architectures. Pretraining with different controller structures could also lead to changes in learned recurrent dynamics, which can alter feedforward controllability without any changes to feedforward architecture; however feedforward controllability spectra were largely similar across architectures, with some reduction in the number of highly feedforward controllable modes for networks with a recurrent controller (Figure 8f).

Next, we implemented learning in these architectures, where adaptation to new decoders was enabled by changes to feedforward weights and to the feedback controller *F*. We observed that for all networks with control bottlenecks i.e. low-dimensional feedback, there was significant variability in learning outcomes, with a similar signature of poorer learning for outside-manifold perturbations (OMPs) (Figure 8h). By contrast, the network with higher-dimensional feedback (Figure 8d) had similarly effective learning of both within- and outside-manifold decoders. Moreover, increasing the number of modifiable parameters, without getting rid of the bottleneck (Figure 8b,c), did not help ameliorate these learning differences. The differential success for outside-manifold decoders across network architectures could be better understood as arising due to differences in controllability, rather than variance observed during the baseline task which was similar across architectures. Thus, adaptive but low-dimensional controls impose strong constraints on short-term learning when effective flowfields in the downstream region are dominated by fixed autonomous (recurrent) dynamics.

## Discussion

Sensory feedback is a key component of skilled motor control and modulates the activity of neurons in primary motor cortex (M1) via direct and indirect pathways [22–25, 69, 70]. In this work, we propose that these feedback-dependent signals may be flexibly updated to “re-steer” neural activity during short-term motor learning. We show how input plasticity, including feedback remapping, is more consistent as a mechanism for fast adaptation to new BCI decoders, with pre-existing input-driven dynamical structure serving as a major constraint for learning. Future work that identifies these dynamics from neural recordings can directly test whether these features predict subsequent learning outcomes.

Interestingly, we find that geometrical features of neural activity, such as the low-dimensional task-specific intrinsic manifold, are insufficient to explain the large variability of learning outcomes. We hypothesize that not only is neural activity constrained within the intrinsic manifold, but that its temporal structure is limited by underlying flowfields. We thus offer a mechanistic hypothesis (learning via input plasticity) and theoretical explanation (low controllability and inconsistent flowfields) for the behavioral and neural observations in previous BCI experiments [14, 66, 71], emphasizing dynamical constraints on learning. Since we identify learning constraints as arising from basic control-theoretic properties of input-driven dynamical systems, rather than from specific aspects of network architecture, these principles can be tested more broadly in other forms of behavioral flexibility.

There are two key features of our framework that distinguish us from related computational studies on motor adaptation [58–62] and that enabled these new insights. First, our use of velocity decoders and inclusion of feedback allowed us to take into account the dynamical nature of neural activity, making the network’s task from open-loop decoding into a closed-loop *control* task. This choice is not only more aligned with the experimental design of BCIs, but allowed us to quantify the ease of producing neural *trajectories*, beyond just neural states. Secondly, we examined the variability between different within-manifold decoder perturbations (WMPs), where all the new decoders were aligned to the intrinsic manifold. Neither the alignment with the original decoders nor the original variance along the decoder (which would be low if the WMP is aligned with trailing PCs) could strongly predict which WMPs would be more learnable. Thus, while geometrical features of neural activity are a useful starting point, they do not fully predict learning outcomes. Rather, we found that constraints on learning arise from the structure of input-driven dynamics that shape both the intrinsic manifold and neural trajectories. Designing more controllable and learnable BCIs thus requires a better understanding of feedback control of cortical dynamics, with more attention to not just the distribution of neural states but the underlying dynamics that govern the time-evolution of those neural states.

### Dynamical constraints on learning

Many experimental and computational studies have demonstrated that collective neural dynamics, i.e. the coordinated and lawful time evolution of neural activity supports many mental and motor computations [1]. These collective dynamics, often characterized by a flowfield, are determined by both the network connectivity and external inputs to the network. While the dynamical systems framework has previously been applied to motor cortex during forelimb-based tasks, we observed that low-dimensional structured dynamics, interacting with task inputs, underlie motor cortical activity in a BCI dataset as well.

Inputs are a key factor for shaping and adapting effective neural dynamics to support behavioral flexibility. Structured recurrent connectivity in motor cortex can be considered as a flexible pattern generator, whose output is modified by initial conditions as well as by external inputs from thalamic and other brain areas [3, 4, 20, 21, 57, 72]. Beyond changes to the attractor landscape, the interaction of recurrent inputs and time-varying control inputs can support new sets of dynamics in nonlinear systems [73]. Such inputs arise naturally in the context of feedback control or ongoing computations in upstream brain regions. Our results suggest that modification of these inputs, either via projection weights or updating nonlinear control policies, can be useful for, and is likely at play during, rapid motor adaptation and other forms of behavioral flexibility. Other recent studies have also demonstrated the potential of learning new inputs to recombine fixed dynamical motifs for fast adaptation in cognitive tasks [10, 13].

However, the success of such adaptation mechanisms depends on the expressivity of the network dynamics and degree of network controllability, both of which are functions of autonomous (recurrent) dynamics as well as input architecture. A recent BCI study observed that primates could not learn to produce reversed neural trajectories within a single session, which would require massive dynamical reorganization on short timescales [71]. Long-term learning of new skills may be less limited by such constraints however, by relying on restructuring of recurrent connectivity. Indeed, multi-day learning of BCI decoders, which is accompanied by reorganization of neural covariance, may rely on such recurrent plasticity [67, 74]. However, given that dynamical structure is relatively constrained during the 1-2 h BCI sessions [71], despite behavioral pressure to change it, similar principles extended to the older BCI learning results [14] are compatible with input plasticity as the basis of learning, as described in our work.

Another key factor for learning was feedback controllability of the BCI decoders. The controllable subspace, a latent property of network and input structure, could be dissociated from a baseline task’s “intrinsic manifold”, and low controllability, rather than low variance per se, explained poorer learning for many outside-manifold decoders. The controllable subspace, and hence the potential of inputs to reshape effective flow along many different directions, can of course be altered by input dimensionality. In our networks, we observed that even more expressive controller networks could not adapt to all decoder perturbations when the control signal was constrained to be low-dimensional. Such low-dimensional signals may arise due to structural connectivity constraints (such as bottlenecks between cerebellum to neocortex), or more likely due to functionally low-rank communication between motor cortex and upstream partners in task-engaged states, as has been characterized in other regions [75, 76]. This constraint on learning due to control bottlenecks arises not just for feedback inputs but for feedforward inputs as well [62].

While the primate BCI studies [14, 71] provide behavioral evidence for “dynamical constraints on neural activity”, our work indicates likely circuit mechanisms as well as a theoretical framework to link controllability in networks to learning speeds. Our work suggests that rapid adaptation is not achieved by “rewiring” the internal connections in motor cortex, but rather by “re-steering” the system through new inputs. This mechanism could implement a fundamental tradeoff between flexibility and stability: it explains why animals can learn certain tasks in minutes without destabilizing existing motor memories, but it also limits what is “learnable” on these fast timescales.

### Modelling feedback for learning in brain-computer interfaces

The use of sensory feedback during BCI use is by now well-established [22, 53, 56, 77–79]. Even simple operant conditioning BCI paradigms generally provide some form of sensory feedback or direct neural stimulation that relates to the network state or task performance (such as a tone modulated by the neuron’s activity), ostensibly to aid task learning [52, 80, 81]. This feedback remains essential for the continued volitional modulation of cortical activity. A recent study using barrel cortex photostimulation as task feedback further found that the structure of the photostimulation pattern determined the rate of learning [51]. Thus, understanding the learning dynamics and representational changes during BCI use requires a deeper consideration of the role of feedback. We observed that altering the feedback mapping could adapt the effective network dynamics, but the extent of this dynamical reorganization determined the learning rate. We similarly expect that learning to use non-intuitive photostimulation patterns requires greater modification of the feedback pathway, and learning may thus be too slow or limited.

As we have currently not included sensory delays, our networks could use immediately available feedback for updating their activity and consequently, the motor output. However, the relatively large delays along the visual pathway (~ 60 ms) necessitate the use of forward internal models that integrate efferent and afferent information to infer the state of the controlled plant, which can guide feedback control. A previous study [77] showed that BCI commands could be best interpreted through the lens of an internal model that combines the last available cursor state and intervening network activity states to predict the current cursor state, adjusting the network output accordingly. Extending our model with delays and explicit forward models will help dissect the network mechanisms that support this computation, and distinguish consequences of updating the internal model versus changing a control policy directly during adaptation.

### Distributed mechanisms for sensorimotor adaptation

What is the neural and synaptic basis of the proposed input plasticity underlying short-term adaptation? Cerebellar circuitry has a well-established role in sensorimotor adaptation [42, 44, 46, 82–88], which likely extends to BCI learning. Sensory prediction errors that arise due to a mismatch between new BCI mappings and previously learnt internal models of the BCI [77] can drive plasticity within the cerebellar cortex, putatively at the granule cell-Purkinje cell synapse, and recalibrate the internal model for both feedforward and feedback control of cortical dynamics [38, 89, 90]. Alternative computational theories also suggest that cerebellar feedback may guide subsequent local plasticity within M1, either by acting as a teaching signal [34, 91] or by estimating surrogate gradients for weight updates [92]. Nonetheless, better characterization of cortico-cerebellar interactions during BCI control as well as learning is a promising next step.

In this study, we used hit rate rather than trajectory loss as a measure of success, similar to quantification of task performance in experimental data. Indeed, in [14], animals were not required to make perfect reaches or stop at the targets, either during the baseline task or the adaptation period. Moreover, sensorimotor adaptation has been shown to have both a reward-driven policy update and an implicit sensorimotor recalibration component [39, 93–97], which are sensitive to target errors and sensory-prediction errors respectively. These learning processes are not entirely independent and the rate of implicit adaptation can be modulated by the reward rate. Explicitly modeling the interaction between the minimization of sensory errors and reward maximization will yield interesting insights into how different learning pressures shape network connectivity and dynamics. Another recent study implicated a parallel form of input plasticity in BCI learning, where low-dimensional *feedforward* inputs are modified, potentially as a cognitive “re-aiming” strategy [62]. While our work supports upstream plasticity as the basis of adaptation, it is possible there are concurrent distributed changes, particularly for longer-timescale learning. Some computational studies have suggested error-or reward-based modulation of local connectivity in M1 during other motor adaptation tasks [34, 98, 99]. Modelling these parallel learning processes and dissecting the multi-region circuits underlying learning is an exciting avenue for future work.

## Methods

### Modelling framework and task description

**Table 1:** Network parameters.

| Parameter | Value | Description |
| --- | --- | --- |
| $N$ | 100 | No. of RNN units |
| $N_{in}$ | 3 | No. of feedforward stimuli (2D target + Go cue) |
| $N_{out}$ | 2 | Output dimensionality (2D velocity) |
| $N_{fbk}$ | 2/4/12 | Feedback dimensionality |
| $\sigma_{in}$ | 0.5 | Initial scaling factor for SD of input weights $W_{in}$ and $W_{fb}$ |
| $\sigma_{rec}$ | 1 | Initial scale for SD of recurrent weights $W_{rec}$ |
| $\sigma_{out}$ | 5 | Scaling factor for SD of readout weights $W_{out}$ |
| $\sigma_b$ | 1 | SD of RNN unit bias term |
| $\tau$ | 5 | Network time constant (dt=1) |
| $\Phi$ | ReLU | RNN unit nonlinearity |
| $\sigma_n$ | 0-0.05 | Standard deviation of activity noise |

#### Task

The network was trained to perform a 2D center-out reaching task, similar to 2D cursor-based BCI tasks, where a readout from network activity is mapped onto the velocity of a “cursor”. On each trial, the network was given a target from 1 of 8 locations. After a Go cue, the cursor was to be moved towards the target, and held at the target thereafter.

#### Network dynamics

The network activity (**r** = ϕ (**x**)) dynamics were defined as:

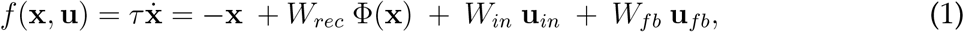

with feedback input set to be the position error (**u**_*fb*_ = **p**\* − **p**). The feedforward input **u**_*in*_ was a 3-dimensional vector that cued the target location **p**\* [cos(*θ*), sin(*θ*)] and the Hold/Go signal as a binary value. *W*_*rec*_, *W*_*in*_, *W*_*fb*_ refer to the recurrent, feedforward input and feedback input weights respectively and the nonlinearity ϕ was chosen to be ReLU unless otherwise noted. The cursor position **p** evolved according to:

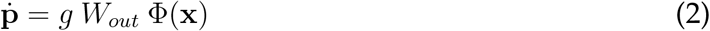

with *W*_*out*_ as the readout weights (“decoder”) and g=1e-3 as a velocity scaling factor.

For simulations where another network *F* was set as the feedback controller, *F* could be a 2-layer feedforward network or a recurrent network. For feedforward controller architectures, the input **y** to the network *F* was a concatenated vector of sampled RNN state (**r**_*s*_ = *M*_*s*_ ϕ (**x**)) and the position error (**p**\* − **p**). This input was then passed via the network *F* (with a chosen architecture) with the final output **x**_*out*_ = *F* (**y**). Sampling the network state **x** enabled the controller *F* to learn a flexible, state-dependent mapping.

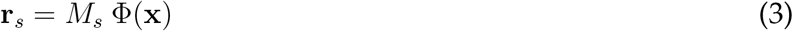

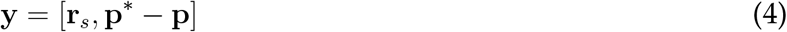

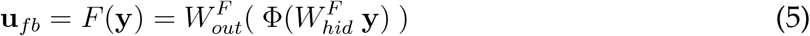

For recurrent controller architectures with hidden state **h**, the input was the error signal, and the recurrent dynamics enabled the controller *F* to use the error trajectory to provide flexible, history-dependent input.

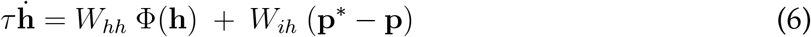

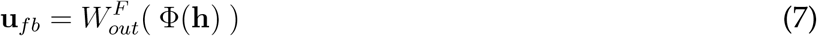

**Table 2:** Feedback controller (*F*) parameters.

| Parameter | Value | Description |
| --- | --- | --- |
| $N_{mf}$ | 10 | Dimensionality of RNN sampling weights $M_s$ |
| $N_{hid}$ | 50/200 | No. of hidden units in F |
| $N_{out}$ | 4/12 | No. of output units in F |
| $\Phi_{hid}$ | 'ReLU' | Nonlinearity at hidden layer |
| $\Phi_{out}$ | None | Nonlinearity at output layer |

#### Training

Weights were initialized from a normal distribution with standard deviation equal to 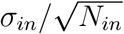 (input weights *W*_*in*_), 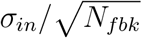 (feedback weights *W*_*fbk*_), 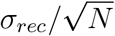 (recurrent weights *W*_*rec*_) and 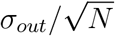 (readout weights *W*_*out*_) respectively. For initial training, the random readout weight matrix *W*_*out*_ was fixed while both recurrent weights (*W*_*rec*_) and input weights (*W*_*in*_, *W*_*fbk*_) were trained via gradient descent (Adam optimizer) with a batch size of 32. For networks with a feedback controller network *F*, the feedback controller was trained instead of *W*_*fbk*_. For all experiments, the readout weights *W*_*out*_ were always kept fixed, either initialized randomly or as decoder perturbations. The minimized loss function was the average squared difference between the desired and actual cursor trajectory, plus weight and activity regularization terms (using the *L*_2_ norm), where the desired trajectory was a sigmoidal function (in time) along the direct vector to the target. During training, we introduced small external perturbations in the form of Gaussian bumps added to the current cursor position (*σ* = 10*dt*, amplitude= 0.02). For testing, we perturbed the cursor in the form of a discrete jump, by moving either the x-or y-coordinate by 0.1. We tested the networks by introducing these perturbations at timesteps 150, 300, and 500.

For experiments with adaptation to decoder perturbations (see later), weights were trained using Adam optimizer with batch size 1. For networks with direct error feedback, either both *W*_*in*_ and *W*_*fbk*_, or only *W*_*rec*_ was trained. For network simulations with controller network *F*, the weights from the hidden to output layer 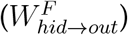 or the recurrent layer (*W*_*hh*_ and *W*_*ih*_) were trained along with *W*_*in*_. Further, the distance to target was penalized during the hold-at-target period (from step no. 300 to 800) but the network was neither constrained to follow a particular trajectory nor required to hold at target until the end of the trial (1500 steps).

#### Weight perturbation

In each training epoch, a pre-perturbation loss (defined above) was computed using the current weights *W* before adding a small perturbation *δW* to each of the trainable weights (independently drawn from a Gaussian distribution). The change in the loss due to this perturbation (*δL*) was used as the error signal, and the weight changes were consolidated as 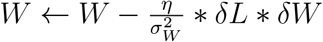.

**Table 3:** Initial training parameters.

| Parameter | Value | Description |
| --- | --- | --- |
| $lr$ | 1e-3/3e-4 | Learning rate (Simple networks/ with controller $F$) |
| $T_{max}$ | 500 | No. of timesteps in a trial |
| $T_{stim}$ | 20 | Stimulus onset timestep |
| $T_{go}$ | [100,200] | Go cue timestep (range) |
| $Ne_{tr}$ | 400-700 | Total number of training epochs (trials) |
| $Ne_{pert}$ | 250-400 | No. of training epochs with cursor perturbations |
| $B_{tr}$ | 32 | Batch size |
| $\alpha$ | $0.03/\sqrt{N}$ | Regularization factor - input weight norm |
| $\gamma$ | $0.03/\sqrt{N}$ | Regularization factor - recurrent weight norm |
| $\beta$ | 0.01 | Regularization factor - Activity norm |

**Table 4:** Altered training parameters for decoder adaptation.

| Parameter | Value | Description |
| --- | --- | --- |
| $lr$ | 1e-3/3e-4 | Learning rate |
| $T_{max}$ | 1500 | No. of timesteps in a trial |
| $Ne_{tr}$ | 200/1500 | Total number of training epochs (trials) |
| $B_{tr}$ | 1 | Batch size |
| $\eta$ | 0.001 | Learning rate for weight perturbation |
| $\sigma_W^2$ | 0.01 | SD of input weight perturbation |

### Decoder manipulations

#### Intrinsic manifold and intuitive decoders

The trained network exhibited low-dimensional activity, allowing us to define an intrinsic manifold using the top 8 principal components (PCs) that captured most of the activity variance. We then performed linear regression between the cursor velocity and activity along these 8 PCs to identify the intuitive decoder *W*_*ID*_ as:

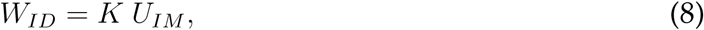

where *K* are the regression coefficients, and the rows of *U*_*IM*_ are the loadings for the top 8 PCs.

#### Within-manifold and outside-manifold perturbations

Following [14], we constructed new perturbed decoders either by permuting the regression coefficients i.e. columns of *K* to construct WMPs, or by permuting columns of *U*_*IM*_ to construct OMPs. To constrain the maximum number of OMPs to be the same as WMPs, we followed the steps in [14], dividing neurons into 8 groups, and permuting grouped columns. (For dividing neurons into groups, we first sorted the neurons by their activity variance, then distributing the top 8 neurons randomly into 8 groups, then the next 8 neurons randomly into those 8 groups, and so on).

We screened all possible WMPs and OMPs and only selected those that passed several criteria, adapted from the conditions in [14]. The mean “open loop velocity” under the WMP/OMP was constrained to be within 0.5x-to-3x of mean velocity observed with using *W*_*ID*_ as readout, and the mean angular difference between the two open loop velocities was less than 90 degrees. The mean closed loop velocity was constrained to be within 0.5x-to-2x of velocity observed with using *W*_*ID*_ as readout. One of these WMPs or OMPs was then set to be the new decoder *W*_*pert*_, and the network weights were retrained to recover task performance.

### Network activity analysis

#### Preparatory and movement PCs

We separately computed principal components of network activity during the delay period (stimulus onset to Go-cue) and during the movement period (Go-cue to reaching target), and computed the principal angles between the subspaces defined by the top 4 PCs for each (referred to as the preparatory and movement subspace).

#### Fixed point analysis

We computed the fixed points of the joint system for different values of feedforward inputs - the target inputs were set to each of the 8 targets, and the Go-cue set to 0 for preparatory epoch and to 1 for the movement/hold-at-target epoch. For example, by freezing the target input to (1,0) and the Go-cue input to 1, we obtain the fixed point during the movement epoch, when the network needs to reach target at (1,0). We obtained the minima of summed neural and cursor speed using gradient descent with an adaptive step size, and confirmed that the minimal speed was less than a small tolerance value. We computed the eigenvalues of the Jacobian at the obtained minima, and confirmed that the real parts were negative, indicative of stable fixed points. For visualization, we concatenated all fixed points during the preparatory (or hold-at-target) epochs, and used the top 3 PCs of this dataset.

#### LDS fitting to model data

We fit an input-driven linear latent dynamical system to the trial-averaged neural activity (for each target) during the movement epoch. Latent dynamics were defined as:

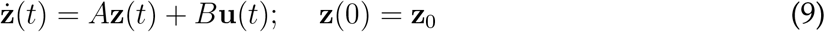

where **z** is the latent variable, and input **u** is a concatenation of the 2D target location and a 2D error signal i.e. vector-to-target from the current position, computed using the ideal trajectory. The neural activity **r** was then defined as a readout of these latent dynamics:

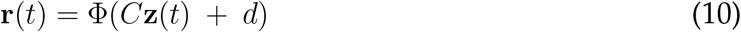

with a ReLU nonlinearity (ϕ) and a bias current *d*.

For model fitting, we first selected a random subset of neurons (~90%) as training set and used the rest for testing. Since we used a non-probabilistic model, we fit the parameters *A, B, C*_*tr*_, *d*_*tr*_, **z**_**0**_ using gradient descent, where *C*_*tr*_ and *d*_*tr*_ corresponds to the rows of *C* and *d* for the training set neurons. The loss was defined as the squared difference between true and fitted neural activity. Next we fixed the latent dynamics (*A, B*, **z**_**0**_) and retrained *C, d* (in practice, *C*_*test*_, *d*_*test*_) on a subset of the targets to identify the observation weights for the test neurons. Model performance with these learnt parameters was then computed as variance explained for the activity of test neurons on the remaining held-out targets. We compared model performance (test *R*^2^) for different numbers of latent variables (k=1 to 14) and typically found k=8 to be the best fit.

#### LDS fitting to BCI dataset

For comparison of network dynamics between experimental recordings and trained computational models, we used an openly available dataset of M1 activity during a center-out BCI reaching task from [17]. This dataset is available at (https://dandiarchive.org/dandiset/000404/0.230605.2024). For analysis we down-sampled data from monkey G by 6x (60Hz to 10Hz), and data from monkey J by 10x (200Hz to 20Hz). We used the binned spike counts as neural observations and centered cursor positions as behavioral observations.

For LDS fits, we did not perform trial averaging as trial lengths were quite variable. Instead, we fit an input-driven LDS to trial-concatenated data using the ssm package (https://github.com/lindermanlab/ssm) [100]. The latent dynamics were again given by equation (4), with the feedback input computed based on the behavioral trajectory on each trial. The probability of observed neural spike counts were based on an inhomogeneous Poisson process with time-varying firing rates given by

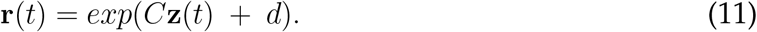

We first selected a subset of the trials (80%) as the training dataset to estimate the model parameters. The variational lower bound for the data log-likelihood (or evidence lower bound, ELBO) was then computed for both the training and test datasets using the estimated parameters. We compared the consistency of the inferred underlying latent dynamics by comparing the eigenvalues of the dynamics matrix *A* across different training subsets. We confirmed that this consistency was not observed in shuffle controls. For one control, activity of each neuron was independently shifted in time, preserving individual neurons’ temporal structure but destroying shared dynamical structure. For the second control, we preserved the population covariance structure, but shuffled the population vector in time. We also identified the rotational frequency and decay timescale of the slowest complex eigenvalue of each fit, and compared it to the inferred rotational modes of the trained models.

For fixed point analysis, we solved the linear equation *A***z** + *B***u** = 0, by setting the feedforward input to each of the 8 targets and the feedback input to be 0 (corresponding to a cursor held stationary at the target). We then computed the first 2 principal components from the set of 8 fixed points, and visualized the inferred latent state 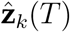 (based on observed neural activity) at the end of each trial *k* in the same 2D space 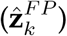. We also used nonlinear classification techniques (K-nearest neighbors) to decode the trial target from this projected latent state 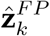, as well as from the full latent state 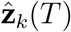, the inferred firing rate 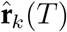, and the true binned spike count at the end of each trial. A similar result was obtained using other nonlinear classifiers.

### Analysis of adaptation to decoder perturbations

#### Task performance

To facilitate comparison with measures of task performance and adaptation defined on experimental data, we computed the hit rate as the fraction of trials where the cursor came within 10% (of total distance) of the target in the defined time window, and used this instead of using the trajectory loss. Similar to experiments in [14], this meant that for successful adaptation, the network was not required to keep the cursor still at the target but just to “hit” the target within a defined time window. The normalized improvement in hit rate was computed as (*h*_*post*_ − *h*_*pre*_)*/*(1 − *h*_*pre*_), where hit rate *h* was computed using performance on 200 trials. This quantified the “amount of learning” and was used as a measure of success of adaptation. We defined the acquisition time for successful (hit) trials as the first timepoint where the cursor crossed this 10% threshold. Progress was defined at each timepoint as the component of the cursor velocity along the current vector-to-target. Behavioral asymmetry was calculated by first computing mean progress (*P*_*i*_) for each of the targets *i* and then computing the index 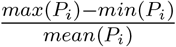.

#### Adaptation learning rate

Individual learning curves were calculated by getting smoothed hit rates across the training trials using a uniform window (of size 20 trials). The smoothing was necessary as hit rates on individual trials took the value of 0 or 1 during this training period. These learning curves were similar across different runs of adapting to the same decoder (different sequence of stimuli). Hence we averaged the learning curves across 3 to 5 adaptation runs to the same decoder and fit a logistic function to these learning curves:

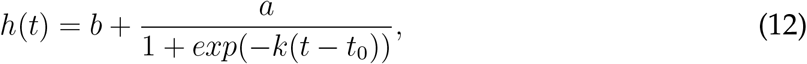

where the rate of learning was quantified as the parameter *k* or the total learning speed (defined as *a* ∗ *k*).

#### Statistical reorganization

To compare similarity of covariance structure between pre- and post-adaptation activity, we took the vectors **s**_*i*_ of relative standard deviation along the top 8 PCs, and computed the similarity as the dot product between the normalized vectors: **ŝ**_*pre*_ ·**ŝ**_*post*_. Similarity of decoders was computed as either the angle between the decoders or the average overlap (inner product) between the corresponding decoder dimensions i.e. for a dimensional decoder, the overlap is 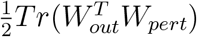.

#### Controllability of readout

To compute controllability properties of the nonlinear system, we can use the notion of Lie brackets [73, 101]. The Lie bracket is a bilinear map on the space of vector fields that captures the non-commutativity of two vector fields *f*_1_ = *F* (*x, u*_1_) and *f*_2_ = *F* (*x, u*_2_):

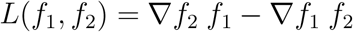

When the Lie bracket is non-zero, it means that the flows do not commute i.e. the order in which the two input-dependent flowfields are applied determines the endpoint. Moreover, the Lie bracket assigns a new tangent vector to each point in the underlying manifold, i.e. varying the input over time to apply *f*_1_ and *f*_2_, in infinitesimal time, produces a new flowfield that differs from the individual input-dependent flowfields. The Lie algebra is the span of all possible vector fields that can be produced by varying the inputs i.e. it is the smallest linear space of vector fields that is closed under the Lie Bracket operation and contains the starting family of fields. For this study we focused on local controllability properties by looking at properties of this vector space at different points along the observed trajectories. Since network dynamics are control-affine in the space of state **x**, i.e. the inputs combine linearly with autonomous dynamics, we computed the Lie algebra corresponding to the family of input-driven vector fields using the iterative method in [73]. This reduced to:

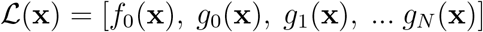

where *f*_0_(**x**) = *W*_*rec*_, *g*_0_(**x**) ≡ *W*_*c*_ = *W*_*in*_ or *W*_*fbk*_, *D*(**x**) = ∇ (ϕ (**x**)) = *diag*(ϕ^′^(**x**)), and

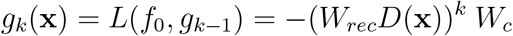

The higher order terms e.g. *L*(*g*_*i*_, *g*_*k*_) reduced to zero as ∇*g*_*i*_ = 0 for *i, k* > 0 with ReLU nonlinearities (all terms included ∇^2^((ϕ (**x**)) = 0).

We examined the spectrum of singular values of ℒ (*x*) for at least 300 different points **x** sampled from neural trajectories under intuitive decoder use. As the spectra were highly similar at different parts of the state space, we averaged them across different sampled states. At each state, we identified the highly controllable subspace *U*_*i*_ as the subspace spanned by the first *i* left singular vectors such that 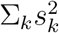 was at least 90% of the total sum (typically 10 modes). To measure controllability of the decoder, we computed the weighted overlap of the highly controllable subspace with the perturbed decoder as 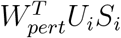, where *S*_*i*_ are the corresponding singular values and normalized it by the overlap between *U*_*i*_ and the intuitive decoder.

#### Changes in flow-fields

The flow i.e. the direction in which the network state **x** would move could be computed at any state along a trajectory (**x, p**) using the network dynamics equation in (1). For networks with controller architecture, the feedback input was computed using the equation (5) or (7). We then computed the L2-norm of the difference in the network flow *f* (or vector-field) along the final observed trajectory, by using pre-training and post-training weights:

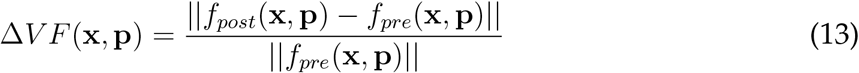

At each state, the flowfield *f*_*post*_ was computed by updating either specific or all weights to the post-retraining values.

#### Regression models

We fit a linear mixed effects model for both the log learning speed and the normalized improvement in hit rate, using different sets of predictors and included a group-dependent (WMP/OMP) intercept.

- *FBK* model: Log feedback controllability of new decoder *W*_*pert*_ (overlap with the top feedback controllable subspace), and log fractional change in feedback-driven vector field
- *G*1 model: Angle between the perturbed decoder and the intrinsic manifold, and change in fractional variance within the original PCs over training
- *G*2 model: Principal angle between perturbed decoder and the original decoder, and initial activity variance along the perturbed decoder
- *G*3 model: Principal angle between perturbed decoder and the original decoder *W*_*out*_, as well as angle with the intuitive decoder *W*_*ID*_

Regression performance was evaluated as the cross-validated explained variance captured on a test set, and recomputed for multiple random partitions (K=40) of the data into training and test sets (80/20).

#### Statistical analysis

Measures are generally reported as median values, along with the 5th-95th percentile values as the confidence interval, unless otherwise noted. For comparison of various measures between WMP and OMP adaptation, we used Mann-Whitney U test. For linear regression, model performance is reported as the median cross-validated variance explained (*R*^2^), and p-values reported based on the F-statistic.

## Code availability

Code to generate the main simulations in the manuscript and to perform data analysis is shared on Github: https://github.com/harshagurnani/FeedbackControlledRNN.

## Data availability

Publicly available datasets [17] used for supplementary analyses are available at (https://dandiarchive.org/dandiset/000404/0.230605.2024).

## Notes

### Competing Interest Statement

The authors have declared no competing interest.

### Summary of Updates

Text changes in introduction/discussion and additional panels in Figs 3-5 to more clearly summarize the conceptual findings.

### Funding

- United States Air Force Office of Scientific Research: FA9550-19-1-0386
- Schmidt Family Foundation

