## Supplementary File for "Feedback control of recurrent circuits imposes dynamical constraints on learning"

### Supplementary Note

#### Dynamics of error-correction

How does the network transform the error signal into a suitable change in the cursor velocity? To understand the dynamics of correction, we analyzed the network in open-loop by clamping the error to be zero except at one timestep. By comparing this network's activity evolution with an unperturbed network, and projecting it through the readout weights $W_{out}$, we could calculate the change in the cursor velocity ($\Delta$Velocity) as a function of time (Figure S1A,B, $\Delta$Velocity is the difference between the perturbed and unperturbed conditions, not between successive timesteps).

What are the possible dynamics of this velocity signal? Since both the input (error feedback) and the output (velocity) are in $\Delta$p coordinates, the optimal solution is for the velocity to change in the direction that is opposite (or anti-parallel) to the error.

- The network could map the feedback input into the readout direction $W_{out}$, and produce a change in velocity that corrects for the error in one or a few timesteps. However, the network would need to produce large activity in the output direction to produce a large velocity output. Since we trained our networks to use this feedback signal to produce a reach trajectory over 100-200 timesteps, and added weight and activity regularization, this solution is unlikely to arise. Moreover, the nonlinear dynamics of the RNN imply a state dependence of the initial $\Delta$Velocity produced by any error signal, suggesting that the initial response is unlikely to have a simple linear additive component.
- The other possibility is that the network produces a nonzero correction of the velocity over many timesteps to slowly correct for the error, with the activity (and velocity) at a reasonable magnitude at all times. The network could again map the error to the output direction such that the initial velocity correction is anti-parallel to the error, and slowly decays over time. In a linear RNN, this would imply that the output direction is an eigenvector of the network dynamics, and the network state evolves unperturbed along any other dimensions (or that the perturbation doesn't propagate for long in other dimensions that feed into the output direction). However, this linear shift to the velocity is still unlikely given the nonlinear dynamics of the RNN. Moreover, by not accounting for the RNN state, the remaining velocity component from the unperturbed dynamics can lead to further changes to the cursor position, potentially leading to a transient increase in the error, or oscillations.
- The third possibility is that the initial response is not aligned with the error, and is small in magnitude so as not to introduce large additional errors. Rather, an anti-parallel component to the velocity slowly emerges as a result of perturbed recurrent dynamics.

We observed that the network's initial response was not aligned with the direction of the error, and that the velocity correction ($\Delta$Velocity) changed in both magnitude and direction over time. The velocity correction grew transiently over ~10-20 timesteps (even though the error was clamped to zero), suggesting that the network's response depends on continued processing by the recurrent network rather than a simple input-output response (Figure S1C). This transient amplification is also a signature of non-normal dynamics. The correction was initially not aligned with the error, but slowly rotated to be in the opposite direction of the error, as shown by the angular difference of $\pi$ between the peak velocity and the initial error (Figure S1D-F). Altogether, we find that rather than a simple transformation of error input into an anti-parallel velocity correction that decays over time, slow recurrent processing shapes the dynamics of correction.

#### Fixed point analyses in feedback-driven networks

Consistent with the identification of stable, decaying dynamics, we observed that neural activity reached stationary states at the end of the reach, suggesting the presence of stable fixed points. However, unlike for position decoders, fixed points of neural activity with velocity decoders may not carry information about the cursor position. For example, neural activity can arrive at the same fixed point via different trajectories in state space, but the final cursor position given by the integration of velocity readout along those trajectories may be different. Similarly, these fixed points may not maintain the cursor at the target location if they produce a non-zero velocity readout. Thus, we reasoned that the robustness to perturbation arises from the presence of stable fixed points of *cursor dynamics* at the target location.

As the dynamics of the RNN and the cursor are coupled, we identified fixed points of the joint RNN-cursor system (i.e locations where the rate of change of network state and cursor state reached a minimum of zero) and confirmed that they were stable attracting fixed points (see Methods). We identified fixed points during both the preparatory and movement (go-to-target) epochs, and separately examined the coordinates corresponding to the RNN and cursor state. For each target, we found unique stable fixed points, and the fixed points changed continuously as we varied the target input (Figure S3a,b). As hypothesized, the corresponding fixed points of the cursor position were either near the centre (during preparatory period) or at the target location (during the go-to-target epoch). Correspondingly, the set of RNN fixed points were arranged in a ring-like structure, mirroring the topology of the targets. This structure of preparatory period fixed points is consistent with analyses of data recorded from primary motor (M1) and dorsal premotor (PMd) cortex during forelimb-based reaches [16, 47, 48].

We repeated this fixed point analysis for a BCI dataset [17] by fitting a latent linear dynamical system to M1 activity recorded during a similar cursor control BCI task in primates (Figure S2). Given the stable M1 dynamics (Figure S2e), we could compute target-specific fixed points of the latent dynamics while the cursor is fixed at the target. Interestingly, the neural activity (or rather, the inferred latent states) at the end of different trials clustered around the corresponding fixed point specific to the reach target, suggesting that a similar strategy may be at play in guiding M1 dynamics during this BCI task (Figure S3c-f). This strategy was not the only possible one to solve the task; alternatively, the network could have used amplifying or unstable dynamics to push the neural state in a target-specific direction, which may potentially be the case during ballistic reaches. Building on previous observations that cortical activity is updated by sensory state during BCI use [22, 77], our simplified model enabled us to propose a dynamical mechanism for how feedback guides neural and behavioral states.

While preparatory fixed points can act as initial states for triggering different trajectories in an autonomous system, separating activity into different regions of the state space also allows input-driven systems to use different local dynamics. More generally, when feedback influences network dynamics, jointly analysing the network and plant (e.g. a cursor or arm) dynamics may be more fruitful or appropriate to understand how these interactions enable “computation” to meet task demands. This is especially applicable to more naturalistic conditions where neural activity controls the kinematics (velocity, force, torques) rather than the position of a plant directly, or where physical forces and bio-mechanical coupling influence plant dynamics.

### Supplementary Figures

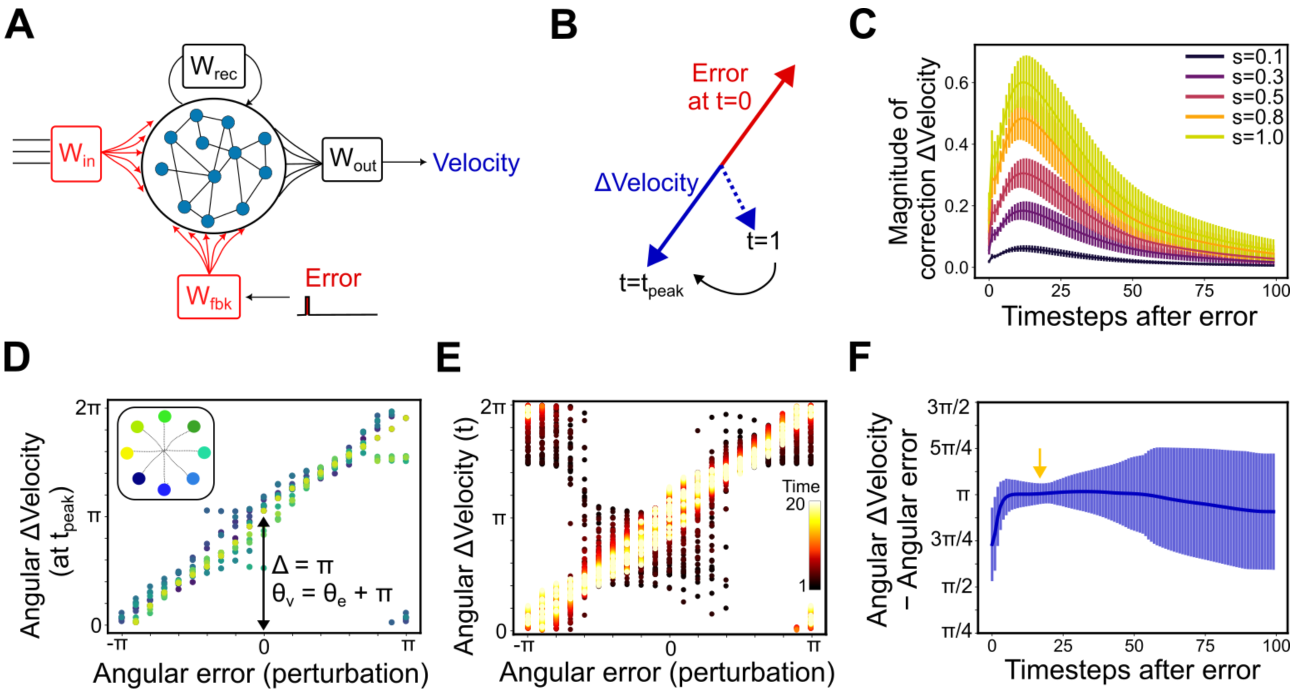


Figure S1: **Slow recurrent processing shapes dynamics of error-correction. (A)** Examining dynamics of error-correction in open-loop (i.e. error-clamp) mode. A single timestep of error is fed to the network via the feedback inputs, and the resultant activity and velocity is compared with that of a zero-error condition. The difference in velocity is shown as ΔVelocity. **(B)** Schematic of how a correction velocity (ΔVelocity) develops after an error (defined by a magnitude $s$ and an angle $\theta_e$). The initial response is small and not aligned with the error. RNN activity slowly evolves such that ΔVelocity is aligned and opposite of the error direction. **(C)** Magnitude of ΔVelocity as a function of time after a single perturbation for different error magnitudes $s$. Thick line and error bars indicate mean and standard deviation across iterations - different reach targets, activity states, and angular errors. **(D-E)** ΔVelocity direction (angle $\theta_v$) as a function of angular error (angle $\theta_e$). **(D)** ΔVelocity direction at the timestep with maximal velocity ($t_{peak}$), colored by different reach targets. **(E)** ΔVelocity direction at different timesteps after perturbation. Different reach targets are plotted separately but colored by the timestep. **(F)** Difference between angular velocity correction and the angular error across different timesteps. Thick line and error bars shows circular mean and standard deviation across iterations of angular errors and reach targets.

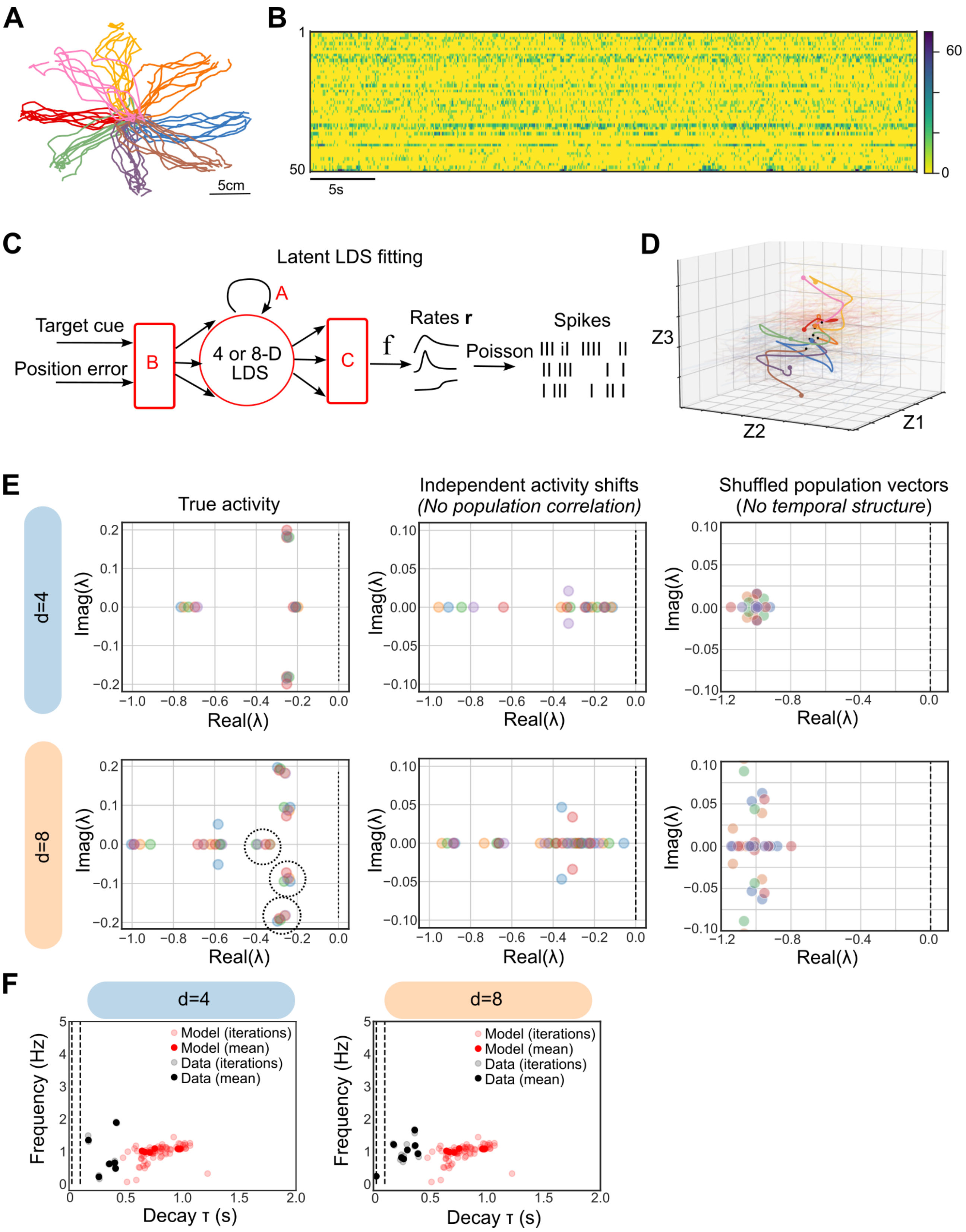


Figure S2: **Latent dynamics identified by LDS fits to primate M1 recordings during BCI-based center-out cursor reaches.**

Figure S2: **Latent dynamics identified by LDS fits to primate M1 recordings during BCI-based center-out cursor reaches. (A)** Cursor trajectories to different targets during an example session. **(B)** Heat map of firing rates of 50 simultaneously recorded neurons. Colorbar indicates firing rate in Hz. **(C)** Schematic of input-driven latent linear dynamical system (LDS) fit to neural data, where neural firing rates were given by a function f (exponential) of a linear combination ($C$) of latent factors, followed by Poisson variability to estimate likelihood of different spike counts. Dimensionality of latent factors (d) was either 4 or 8. **(D)** Latent trajectories from an example session. Bold lines indicate target-specific mean trajectory (color scheme same as A), black dots indicate initial state, and larger colored dots indicate end states. **(E)** Consistency of eigenvalues of estimated dynamics matrix $A$ across different training sets each consisting of different subsets of trials (different colored scatter), for true data (left), circularly shifted activity (independently shifted across neurons) (middle), and shuffled population vectors (right). Top row shows latent LDS fits with d=4 factors, bottom row shows latent LDS fits with d=8 factors. Dotted circles shows cluster of consistent eigenvalues. **(F)** Rotation frequency and decay timescale of the slowest rotational mode for primate M1 data (black; n=8 sessions, N=2 animals), for fits with d=4 (left) and d=8 (right), as well as for trained RNNs (n=3 networks) used in the main paper (red) with d=10 latent factors. Light circles show values estimated from different fitting iterations (different training trials), dark circles show the mean for individual datasets or networks.

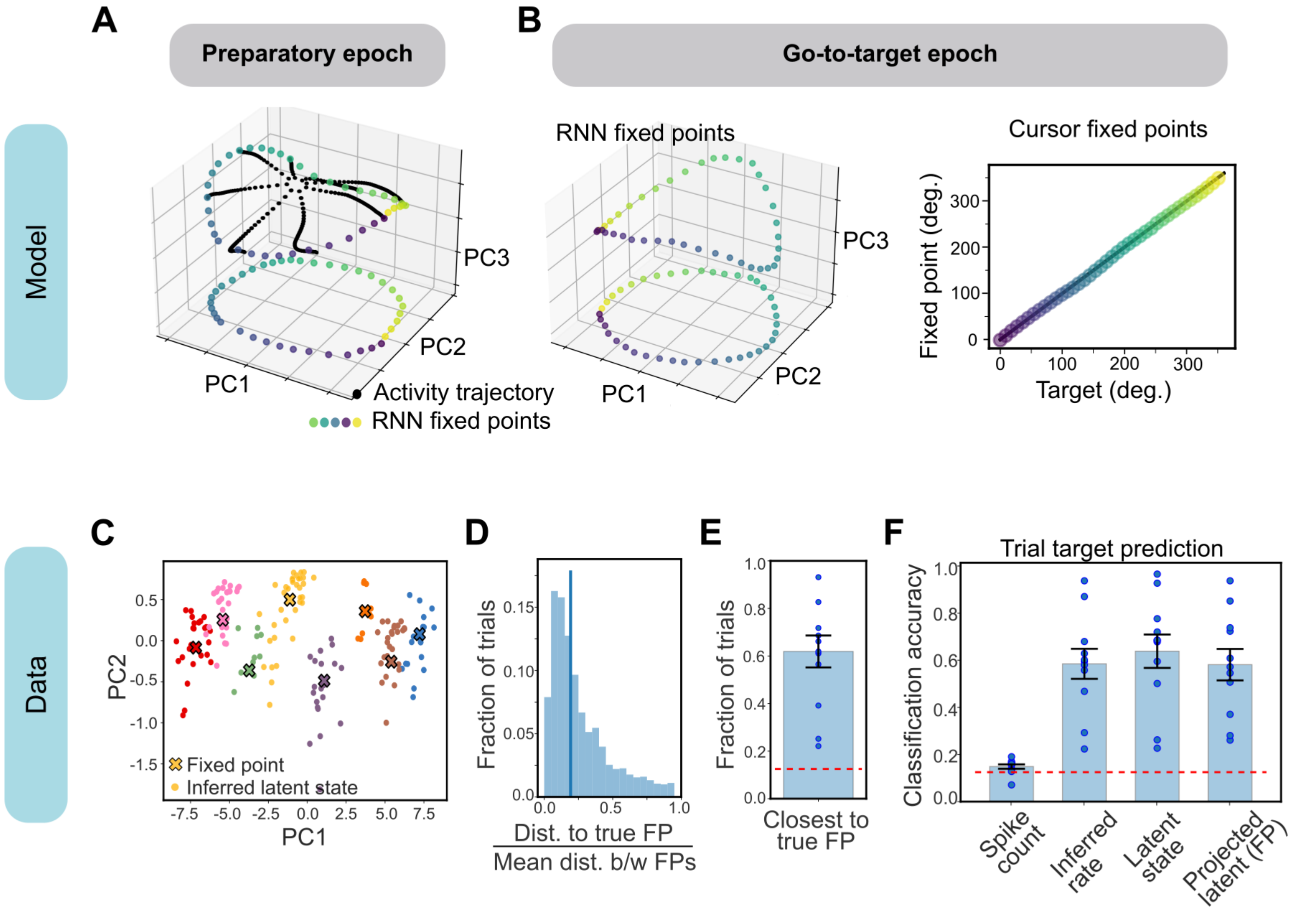


Figure S3: **Fixed points of input-driven dynamics. (A-B)** Fixed points of joint dynamics in the model. **(A)** RNN-coordinates of target-specific fixed points during the preparatory period (Go-cue=0), plotted in the top 3 PC space, along with their projection in the PC1-PC2 plane. 8 example preparatory trajectories to these fixed points are denoted in black. **(B)** (Left) RNN-coordinates of target-specific fixed points during the movement period (Go-cue=1), in the space of top 3 PCs, as well as their projection in the PC1-PC2 plane. (Right) Angular position of the cursor fixed point versus the true target. The target location becomes a stable fixed point, supporting error-correction properties. **(C-F)** Fixed points for the low-dimensional latent dynamics fit to BCI data (same as Figure S2). **(C)** Target-specific fixed points (denoted by crosses), plotted in the top 2 PC space, computed with feedback input set to zero (cursor assumed to be at target) for an example session The inferred latent state at the end of each trial (denoted by circles) is projected onto this space, colored by trial target. **(D)** Distribution of normalized distance of the last latent state on each trial with the correct target-specific fixed point (n=11 sessions). The normalization factor is the mean distance between different fixed points. **(E)** Fraction of trials (per session) where the last inferred latent state was closest to the correct target-specific fixed point (n=11 sessions). **(F)** Nearest-neighbor classification accuracy of the trial target from either raw spike counts, inferred firing rate (denoised), inferred latent state, or latent state projected into the top 3 PCs of fixed points, all taken at the last timestep of each trial. Dashed red line denotes chance accuracy. For (E) and (F), bar and errorbar indicate mean and standard deviation, scatter denotes individual sessions.

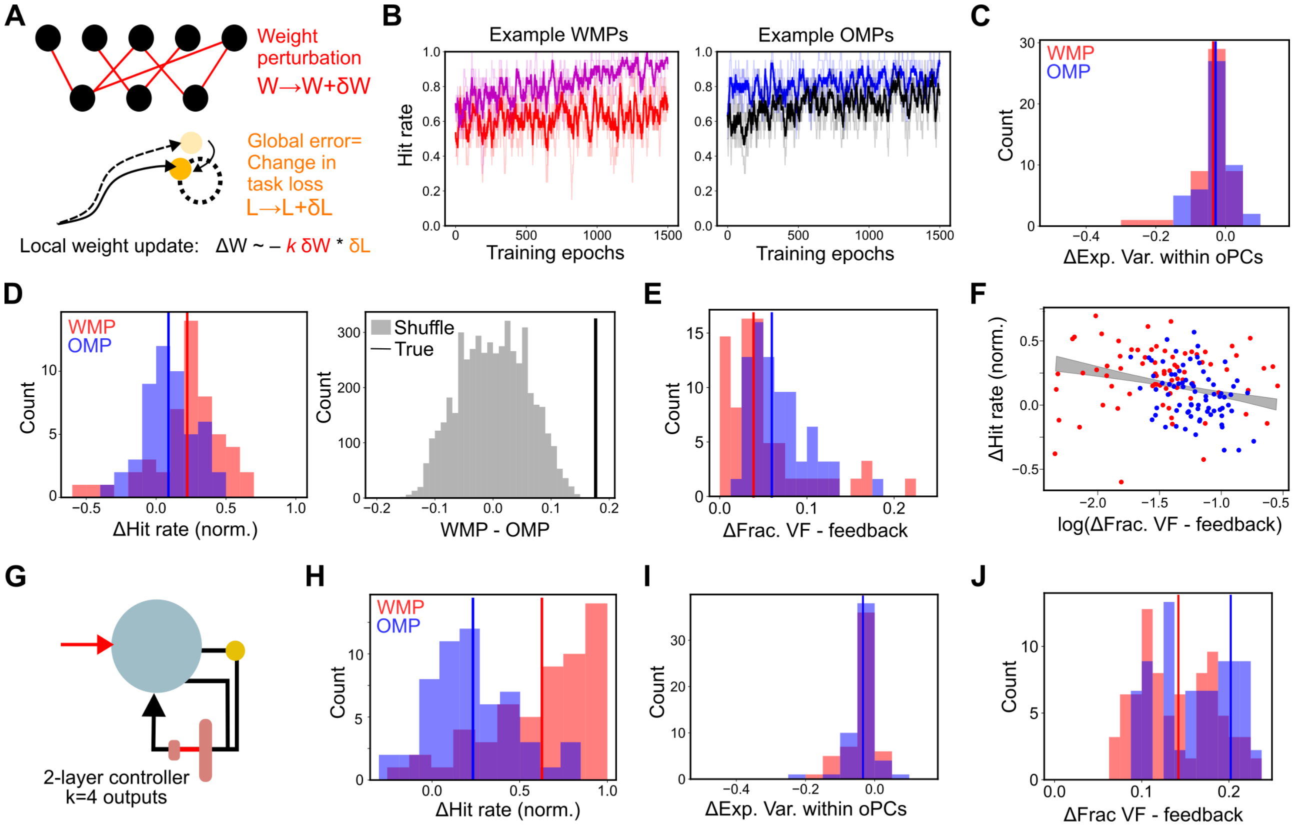


Figure S4: **Adaptation using weight perturbation. (A)** Schematic of weight perturbation as a local learning rule for adapting to decoder perturbation. At each timestep, the change in task loss due to a small, independent perturbation at each weight is measured as the global error signal, and the weights are updated either in or away from the direction of the perturbation, depending on the whether the loss decreased or increased. **(B)** Example learning trajectories (change in hit rate over training epochs) for two WMPs and two OMPs. **(C)** Distribution of change in fractional variance within the original PC space (intrinsic manifold), for both WMPs (red) and OMPs (blue). **(D)** (Left) Distribution of normalized improvement in hit rate for WMPs and OMPs. (Right) Difference between median WMP and OMP learning, for shuffled distributions (gray) and true data (black line). True difference between WMPs and OMPs is significantly larger than shuffle control (>99th percentile). **(E)** Distribution of change in feedback-driven flowfields. **(F)** Normalized change in hit rate versus change in feedback driven flowfields. Gray shaded area shows confidence interval of linear fits. **(G-J)** Learning via weight perturbation for network with 2-layer controller $F$ with 4-dimensional controller output. **(H)** Distribution of change in hit rate for WMPs and OMPs. **(I)** Distribution of change in fractional variance within original PC-space (intrinsic manifold). **(J)** Distribution of change in feedback-driven flowfields.

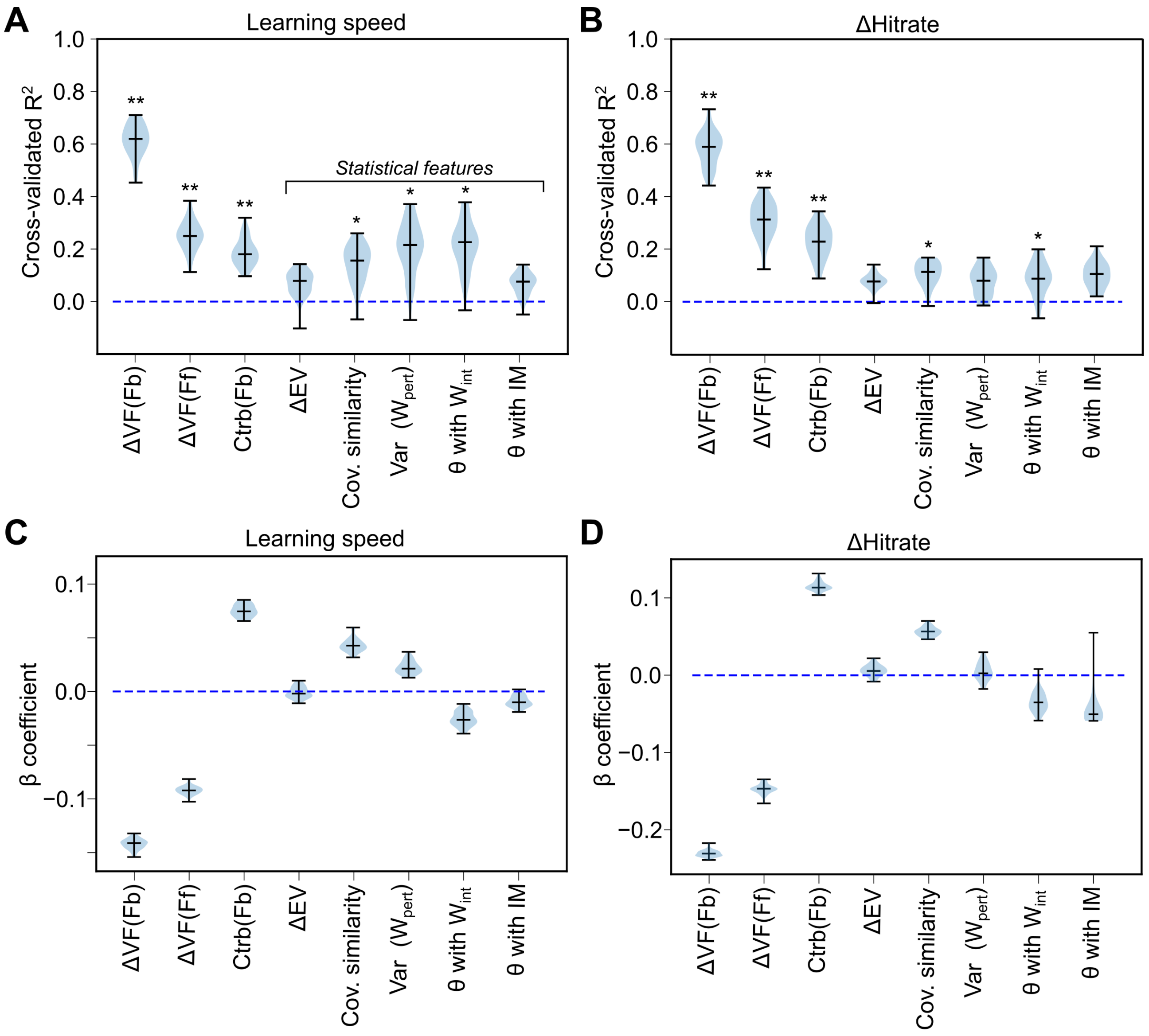


Figure S5: **Regression models for predicting learning variability. (A-B)** Cross-validated $R^2$ of linear mixed-effect model for predicting (A) learning speed and (B) normalized change in hit rate using different predictors (distribution over k=40 fitting iterations, fit to total n=305 perturbations). ** indicates predictors were highly significant ($p<1e-3$), * indicates $1e-3<p<5e-2$. **(C-D)** $\beta$ coefficient indicating strength of association between each predictor and (C) learning speed and (D) change in hit rate.

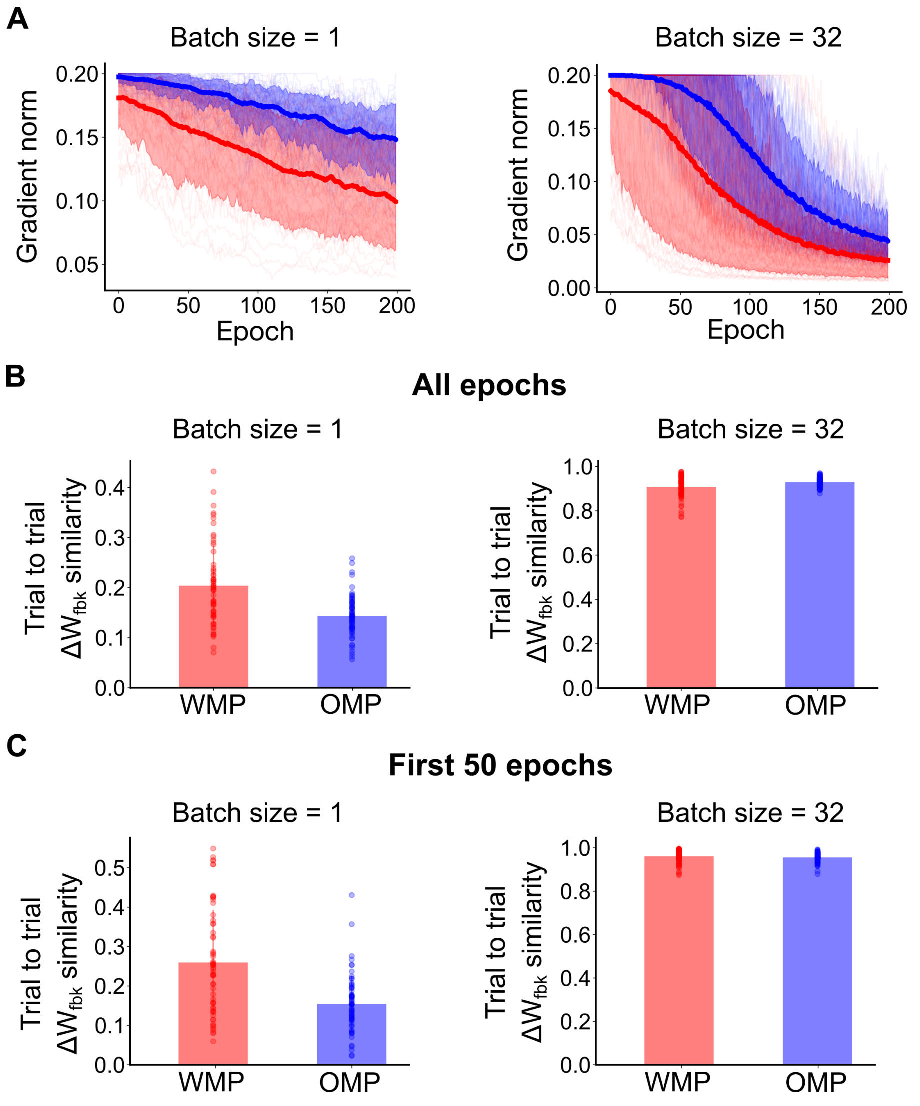


Figure S6: **Gradient estimates during adaptation to WMPs and OMPs for varying batch sizes.** Magnitude of estimated gradients and similarity of weight updates, for retraining using batch size of 1 (left) or batch size 32 (right). **(A)** Gradient magnitude across training epochs. Light lines indicate individual decoder perturbations, thick lines and shaded area indicate median and 5-95th quantile. **(B-C)** Cosine similarity between successive feedback weight updates for WMPs and OMPs, averaged across all epochs in (B), or only for the first 50 epochs in (C). Scatter denotes different decoder perturbations, bars indicate mean across perturbations.

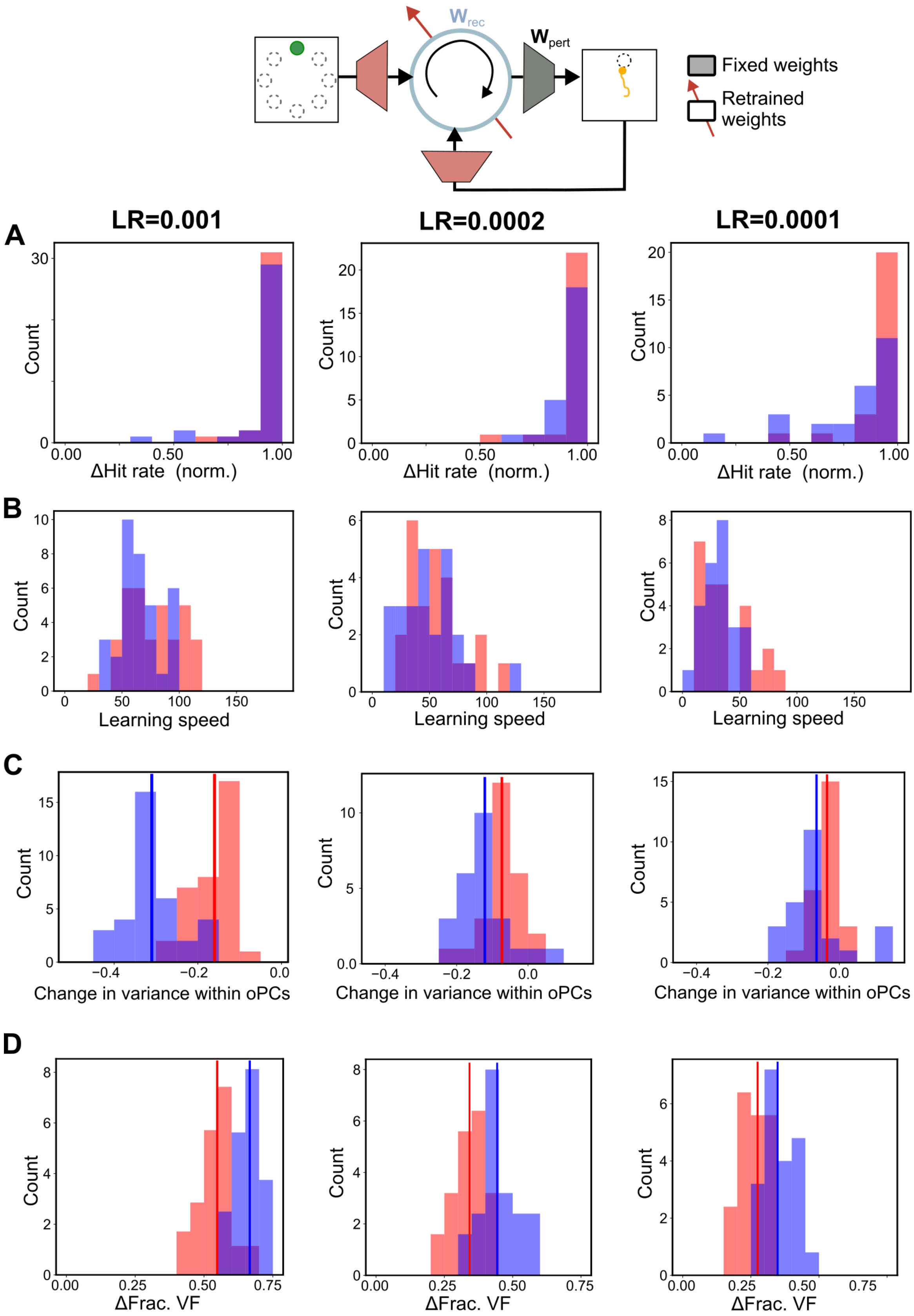


Figure S7: **Adaptation via recurrent plasticity with different learning rates.**

Figure S7: **Adaptation via recurrent plasticity with different learning rates.** Distribution of **(A)** normalized change in hit rates, **(B)** estimated learning speeds (from logistic fit to individual learning curves), **(C)** change in fractional variance with original PCs, and **(D)** normalized change in total flow, for various WMPs (red) and OMPs (blue), using different learning rate parameters for the ADAM optimizer.
